# PEDF-Rpsa-Itga6 signaling regulates cortical neuronal morphogenesis

**DOI:** 10.1101/2020.01.06.895672

**Authors:** Sara M. Blazejewski, Sarah A. Bennison, Ngoc T. Ha, Xiaonan Liu, Trevor H. Smith, Kimberly J. Dougherty, Kazuhito Toyo-oka

**Author notes:** **Corresponding Author** Kazuhito Toyo-oka, Ph.D., Department of Neurobiology and Anatomy, Drexel University College of Medicine, 2900 W Queen Lane, Room 186, Philadelphia, PA 19129 USA.

## Abstract

Neuromorphological defects underlie neurodevelopmental disorders and functional defects. We identified a function for ribosomal protein SA (Rpsa) in regulating neuromorphogenesis using *in utero* electroporation to knockdown Rpsa, which results in apical dendrite misorientation, fewer/shorter extensions with decreased arborization, and decreased spine density with altered spine morphology. We investigated Rpsa’s ligand, pigment epithelium-derived factor (PEDF), and interacting partner on the plasma membrane, Integrin subunit α6 (Itga6). Rpsa, PEDF, and Itga6 knockdown cause similar phenotypes, with Rpsa and Itga6 overexpression rescuing morphological defects in PEDF deficient neurons *in vivo*. Additionally, Itga6 overexpression increases and stabilizes Rpsa expression on the plasma membrane by preventing ubiquitination of Rpsa. GCaMP6s was used to functionally analyze Rpsa knockdown via *ex vivo* calcium imaging. Rpsa deficient neurons showed less fluctuation in fluorescence intensity, suggesting defective sub-threshold calcium signaling. Our study identifies a role for PEDF-Rpsa-Itga6 signaling in neuromorphogenesis, thus implicating these molecules in the etiology of neurodevelopmental disorders and identifying them as potential therapeutic candidates.

## Introduction

Neuronal morphogenesis transforms an immature spherical neuron into a mature neuron with a complex structure. Investigation of the mechanisms that drive neuromorphogenesis has clear applications for improving treatments for neurodevelopmental disorders, as improper neurite formation and dendritic development have been strongly associated with mental retardation disorders and autism spectrum disorder (1–5). Inappropriate dendritic arborization may also impact inputs and signaling efficiency between the synapse and soma (6, 7). Cortical pyramidal neurons have one apical dendrite, which typically extends directly towards the cortical plate. Apical dendrite orientation is crucial in determining synaptic connectivity and is important for neuronal function, as misorientation of the apical dendrite could cause formation of aberrant connections (8). The density and morphology of dendritic spines is another important aspect of neuromorphogenesis. Dendritic spines are the site of over 90% of excitatory synapses in the central nervous system (9). Spine morphology directly relates to synapse function and altered spine phenotypes can result from neurodevelopmental disorders (10, 11).

Ribosomal protein SA (Rpsa), also known as the 67-kDa laminin receptor, functions in a variety of roles including cell anchoring via laminins, ribosomal biogenesis, and chromatin and histone binding (12). Mature Rpsa is embedded in the plasma membrane and may be concentrated in lipid rafts, suggesting it functions as part of a signaling complex (13). Rpsa’s role as a laminin receptor has been extensively studied, revealing its importance in interacting with the extracellular matrix (12, 14–16). Additionally, Rpsa initiates signaling for protection against cell death induced by serum withdrawal in Neuroscreen-1 cells upon treatment with laminin-1, demonstrating an important role for Rpsa in a neuron-like cell line (17). However, the specific role of Rpsa in cortical development remains unknown.

Rpsa binds the secreted glycoprotein pigment epithelium-derived factor (PEDF) (18). PEDF has neurotrophic properties and protects neurons in a variety of regions throughout the central nervous system against excitotoxicity and oxidative damage (19). Recently, PEDF has been shown to promote axon regeneration and functional recovery in dorsal root ganglion neurons after spinal cord injury (20). PEDF expression declines with age and is downregulated by more than 100-fold in aged human fibroblast as compared to young human fibroblast, suggesting an important role for PEDF in early development (21). Additionally, *Serpinf1*, coding for the PEDF protein, is encoded in a clinically relevant region of chromosome 17p13.3 known as the Miller-Dieker Syndrome critical region that is frequently deleted or duplicated in a variety of neurodevelopmental disorders (22, 23). Thus, the relationship between PEDF and its receptor Rpsa in the developing cortex is of interest.

Binding of PEDF to Rpsa may involve a receptor complex that includes the integrin subunit α6 (Itga6), which binds to the β4 subunit to form a complete integrin. Co-localization of Rpsa and Itga6 on the plasma membrane may result from initial co-localization of Rpsa and Itga6 in cytoplasmic complexes, leading to trafficking to the membrane together (12, 24). Furthermore, Rpsa and Itga6 expression may be co-regulated (25). The Rpsa-Itga6 complex is hypothesized to have a role in regulating or stabilizing Rpsa’s interaction with laminin, suggesting that an interaction between Rpsa and Itga6 could be important for other signaling mechanisms like PEDF-Rpsa signaling (25).

We analyzed the functions of Rpsa in cortical neuronal development by performing *in utero* electroporation at E15.5 to induce knockdown (KD) of Rpsa in layer 2/3 cortical pyramidal neurons (26–28). We found that Rpsa deficient cells show defects in apical dendrite orientation, initiation and elongation of dendrites, dendritic branching, and dendritic spine density and morphology *in vivo*. Similar defects are observed following Rpsa KD, PEDF KD, and Itga6 KD. Rpsa overexpression (OE) rescued morphological defects resulting from PEDF KD *in vivo*, suggesting that PEDF initiates Rpsa signaling to regulate neuromorphogenesis. Itga6 OE rescued dendrite formation deficits caused by PEDF KD. Itga6 OE also increases and stabilizes Rpsa expression on the plasma membrane by preventing ubiquitination of Rpsa, thus indicating an important role for Itga6 in this signaling mechanism. Additionally, we show that morphological changes associated with Rpsa KD impact sub-threshold calcium activity. Our study identifies functions of the PEDF-Rpsa-Itga6 signaling pathway in cortical neuromorphogenesis and implicates this signaling pathway in the etiology of neurodevelopmental disorders.

## Results

### Expression of Rpsa in the Developing Cerebral Cortex

The expression pattern of Rpsa has not been previously investigated in the developing brain. Therefore, it was unknown at which levels or developmental time-points Rpsa is expressed, which should be established when investigating the function of Rpsa during development. To quantify the expression of Rpsa protein levels in the developing cerebral cortex, we performed Western blots using lysate of the cerebral cortex from embryonic day (E) 13.5 to P7 samples (Fig. 1A and B). The Rpsa protein is moderately expressed in the developing cortex during early embryonic stages (E13.5) and then expression decreases but persists to at least P7. This suggests that Rpsa may be crucial at earlier embryonic stages. We also analyzed the spatial distribution of the Rpsa protein by immunofluorescence using E18.5 brain sections (Fig. 1C). Rpsa is expressed throughout all cortical layers and in the cytoplasm, but not the nucleus at E18.5. Furthermore, staining of primary cortical neurons confirmed the expression of Rpsa in neurons by staining with the neuronal marker, class III β-tubulin, showing that Rpsa is highly concentrated in the soma and proximal extensions, but is not found in the nucleus or more distal extensions (Fig. 1D). Therefore, functions of Rpsa that can be associated with these cellular localizations, such as interacting with the extracellular matrix, are likely more important at this time-point.

**Figure 1.**
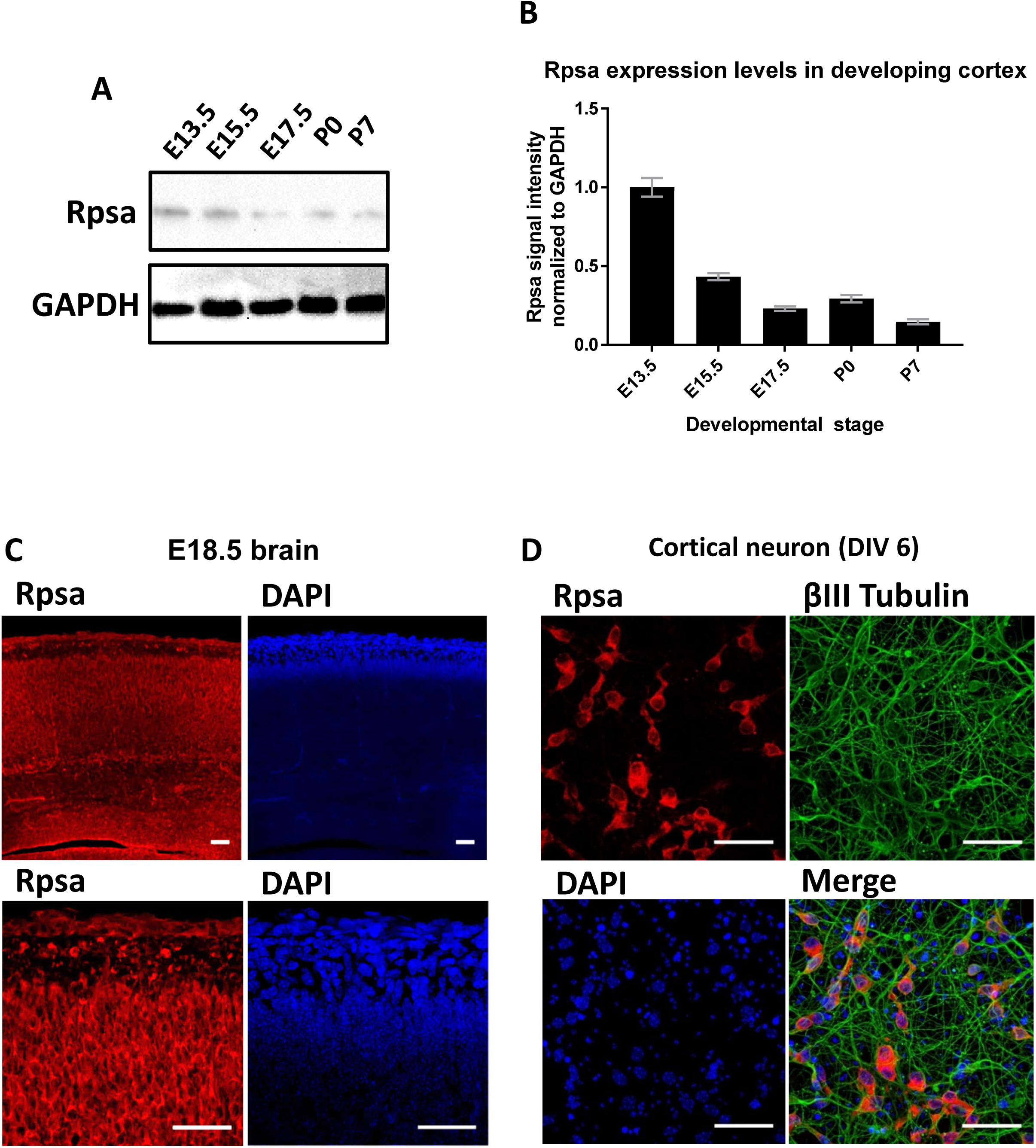
Expression of Rpsa in the developing cerebral cortex and in primary cortical neuronal culture. A) Western blot showing Rpsa expression level in lysates of the cortex at various developmental time-points. B) Quantification of the Western blot showing Rpsa expression levels in the cortex done using ImageJ. Three biological/technical replicates were used (n = 3). Band intensity was quantified and normalized to glyceraldehyde 3-phosphate dehydrogenase (GAPDH). C) Immunofluorescence staining of wild-type brain slices at E18.5 showing the spatial distribution of Rpsa expression (top x20, bottom x63). Scale bars = 50 µm. D) Immunofluorescence staining of cortical primary neurons after 6 days in culture. Staining shows distribution of Rpsa and neurons are double-positive for neuronal marker Tuj1 (βIII tubulin).

### Rpsa KD is associated with apical dendrite misorientation at P3 that can be rescued by Rpsa OE

KD efficiency of our Rpsa CRISPR construct was tested via Western Blot and the knockdown efficiency was approximately 77% (Suppl. Fig. 1A and B). Genomic alteration by Rpsa CRISPR was mainly accomplished by deletions, but cases of single nucleotide additions were also observed (Suppl. Fig. 1 C and D). To further confirm Rpsa KD, we performed immunostaining for Rpsa in primary cortical mouse neurons that were transfected with Rpsa CRISPR. Neurons positive for Rpsa CRISPR (43,881.13 CTFC ± 5,156.70) had significantly decreased fluorescence intensity for Rpsa, as compared with untransfected controls (85,504.51 CTFC ± 9,189.28) (Suppl. Fig. 1 E and F).

To investigate apical dendrite orientation, we performed *in utero* electroporation at E15.5 to induce a CRISPR/Cas9 mediated KD of Rpsa in layer 2/3 cortical pyramidal neurons. The orientation of the apical dendrite is an important feature of neuronal morphology, since the angle at which the apical dendrite extends with respect to the cortical plate is important for neuronal connectivity (8, 29, 30). Conducting this analysis at P3 allowed for easy identification of the apical dendrite, since neuronal morphology is relatively simple at this stage. We found that Rpsa KD cells (34.2° ± 3.61) had a significantly greater mean angle from the soma than the control CRISPR cells (14.1° ± 1.63) (Fig. 2A and B). The frequencies for degree of angle in Rpsa KD cells were more broadly distributed across a wider range from −30° to 30°, while angle measurements were closer to 0° and more narrowly distributed in control CRISPR cells (Fig. 2C). This suggests that Rpsa deficient cells have apical dendrites that are inappropriately positioned at an abnormal angle, which may lead to formation of aberrant synapses. The Rpsa KD phenotype could be rescued by CRISPR-resistant wild-type Rpsa OE (11.8° ± 1.13), resulting in a phenotype comparable to the control CRISPR + control OE group (11.7° ± 1.25) (Fig. 2). The possibility of the phenotype being caused by an off-target effect of the CRISPR is minimal, since the Rpsa KD phenotype was able to be rescued by CRISPR-resistant Rpsa OE.

**Figure 2.**
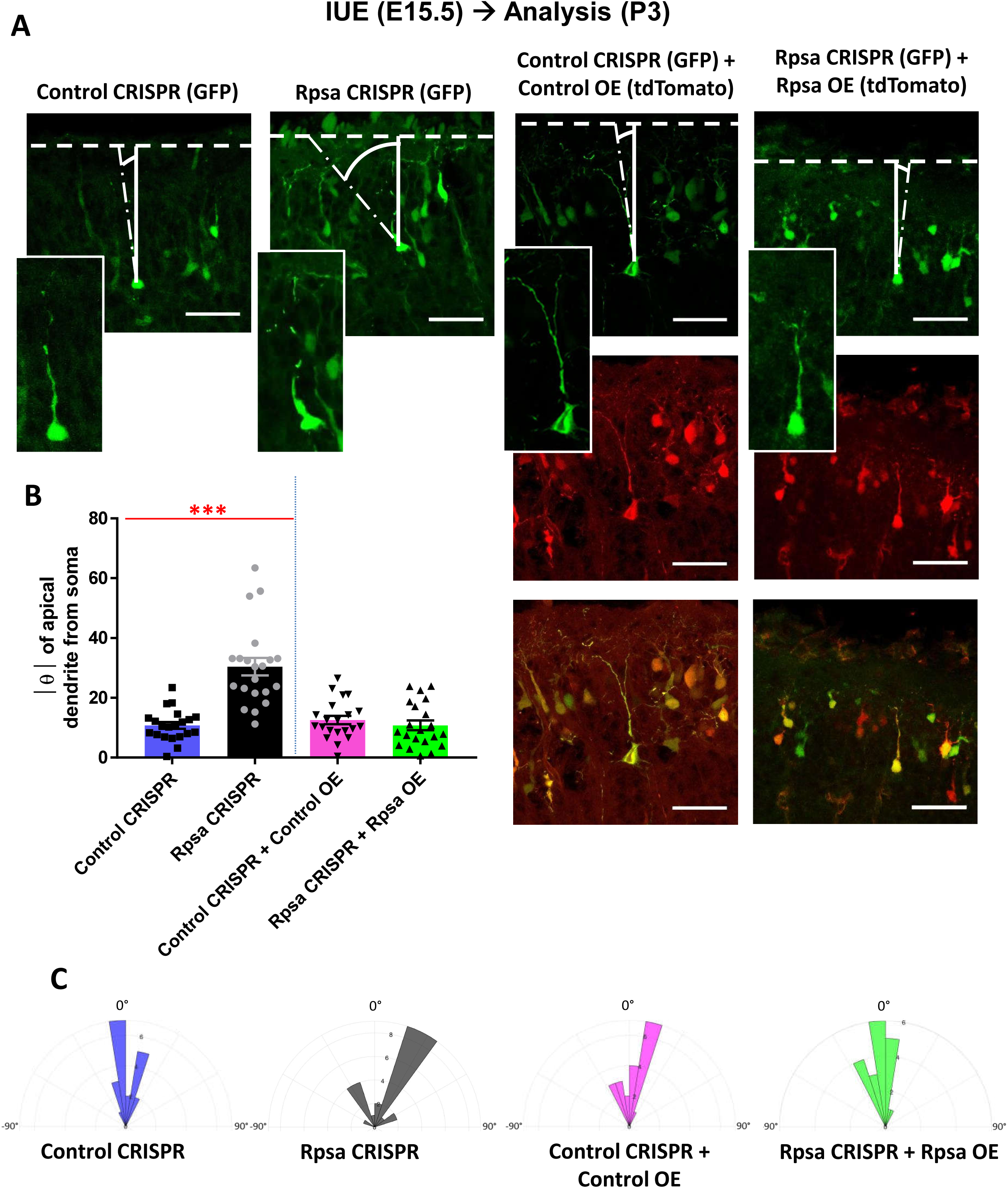
Rpsa CRISPR is associated with apical dendrite misorientation at P3 that can be rescued by Rpsa OE. A) Neurons transfected via *in utero* electroporation at E15.5, orientation of the apical dendrite measured using the angle at which the apical dendrite extends from the middle of the soma relative to the cortical plate, scale bar = 50 µm B) One-way ANOVA showed a significant difference in mean angle of apical dendrite from soma (F(3,76) = 22.620, *p* < 0.0005). Tukey post hoc test showed degree of angle of apical dendrite from middle of soma is significantly greater for the Rpsa CRISPR group (30.4° ± 3.09) compared to control CRISPR (10.7° ± 1.20) (*p* < 0.0005). There was no significant difference between Rpsa CRISPR + Rpsa OE (10.8° ± 1.71) and control CRISPR + control OE groups (12.5° ± 1.49) (*p*=0.924). For all groups n=21 cells from 3 mice. Data are represented as mean ± SEM. C) Dendrograms show frequency distribution of angle measurement.

Additionally, we measured the length of the apical dendrite and found no significant difference between Rpsa KD (27.7 µm ± 1.4) and control CRISPR (27.5 µm ± 1.6) (Suppl. Fig. 2). This suggests that there is no defect in the early period of dendrite elongation.

### Rpsa KD is associated with defects in neuronal morphogenesis at P15 that can be rescued by Rpsa OE

A decrease in the number of dendrites extending from the soma suggests that defective dendrite initiation has occurred, while a decrease in dendrite length suggests a defect in elongation of dendrites. These early pivotal processes have implications for the remaining stages of neuronal morphogenesis, such as dendritic arborization. *In utero* electroporation was used to induce a CRISPR/Cas9 mediated KD of Rpsa at E15.5 and neuronal morphology was analyzed at P15, since neuritogenesis is well-established by this time point. All Rpsa deficient neurons reached the cortical plate (CP) by P15, but the somas were more broadly distributed (Suppl. Fig. 3). The Rpsa KD neurons had significantly fewer and shorter extensions with less complex branching as compared with the control CRISPR neurons (Fig. 3A). The mean number of dendrites from the soma in Rpsa deficient cells was 2.9 ± 0.24, while the mean number of dendrites, including apical and basal dendrites, from the soma of control neurons was 4.2 ± 0.24 (Fig. 3B). The mean dendrite length in Rpsa deficient cells was 20.9 µm ± 2.0 as compared to control neurons with a mean dendrite length of 49.5 µm ± 2.2 (Fig. 3C). Additionally, Sholl analysis of Rpsa deficient neurons indicates that the Rpsa KD neurons have a statistically significant defect in dendritic branching at radial distances 20-75 µm (Fig. 3D). The Sholl profile is consistent with our analysis of dendrite number and length, since it also indicates that Rpsa deficient cells have fewer and shorter dendrites than controls. All described morphological deficits from Rpsa KD were rescued by CRISPR-resistant Rpsa OE, making it unlikely that the observed phenotype was due to off-target effects (Fig. 3). This suggests that Rpsa signaling is crucial in regulating dendrite formation, during both initiation and elongation, in addition to dendritic arborization. Since the apical dendrite was correctly extended as observed at P3 (Suppl. Fig. 2), the defects in the extension of dendrites are mainly in basal dendrites.

**Figure 3.**
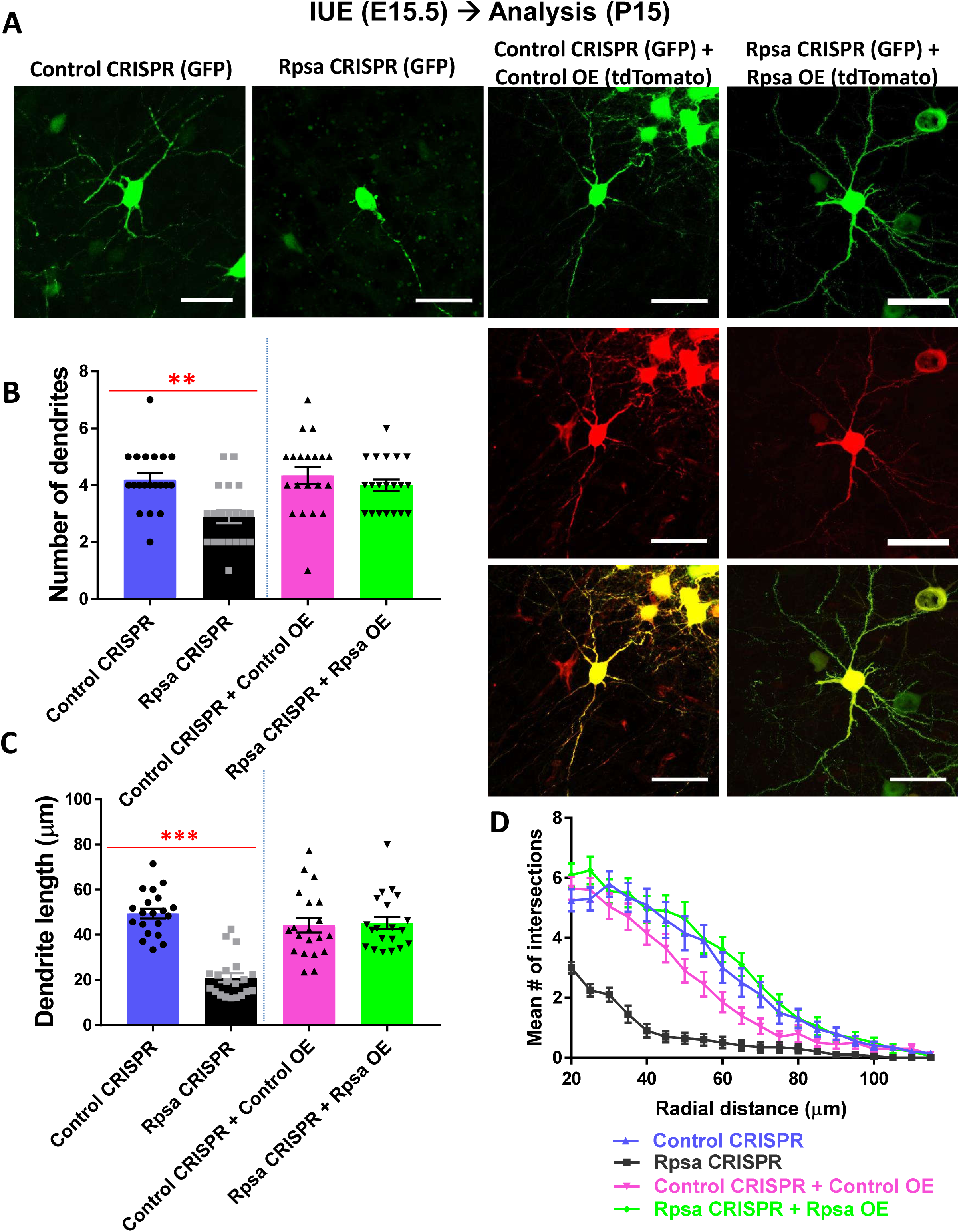
Rpsa KD is associated with defects in neuronal morphogenesis at P15 that can be rescued by Rpsa OE. A) Neurons transfected via *in utero* electroporation at E15.5, scale bars = 50 µm B) One-way ANOVA shows a significant difference (F(3,76) = 7.024, *p* < 0.0005). Tukey post hoc test shows the mean number of dendrites is significantly lower for Rpsa CRISPR group (2.9 ± 0.240,) as compared to control CRISPR (4.2 ± 0.236) (*p*=0.002). There was no significant difference between Rpsa CRISPR + Rpsa OE (4.0 ± 0.205) and control CRISPR + control tdTomato OE groups (4.3 ± 0.301) (*p*= 0.751). For all groups n=20 cells from 3 mice. Data are represented as mean ± SEM. C) One-way ANOVA shows significant difference (F(3,76) = 24.361, *p* < 0.0005). Tukey post hoc test shows dendrite length is significantly lower for Rpsa CRISPR group (20.90 ± 2.04) as compared to control CRISPR (49.51 ± 2.19) (*p*<0.0005). There was no significant difference between Rpsa CRISPR + Rpsa OE (45.29 ± 2.75) and control CRISPR + control tdTomato OE groups (39.99 ± 3.31) (*p*=0.992). For all groups n=20 cells from 3 mice. Data are represented as mean ± SEM. D) Sholl analysis shows a decrease in branching complexity in Rpsa deficient cells that can be rescued by Rpsa OE. Two-way ANOVA with Dunnet multiple comparison test shows a significant decrease in branching for the Rpsa CRISPR group at radial distances 20-75 µm as compared to the control CRISPR group, F(57,1520) = 5.593, *p*<0.0001. For all groups n=20 cells from 3 mice.

### Rpsa and PEDF show similar defects in neuronal morphology after KD and Rpsa OE can rescue morphological defects after PEDF KD

Since PEDF is a known ligand of Rpsa that has neurotrophic properties and is encoded in the clinically relevant Miller-Dieker Syndrome critical region of chromosome 17p13.3, we investigated whether PEDF binding was responsible for initiating Rpsa signaling to regulate neuronal morphogenesis. We used PEDF shRNA in combination with *in utero* electroporation to conduct an initial analysis of PEDF deficient neurons *in vivo*. KD efficiency of the PEDF shRNA was tested via Western Blot and the knockdown efficiency was approximately 84% (Suppl. Fig. 1A and B). Also, we performed immunostaining for PEDF in primary cortical neurons that were transfected with PEDF shRNA. Neurons positive for PEDF shRNA (11,735.13 CTFC ± 2,019.42) had significantly decreased fluorescence intensity for PEDF, as compared with untransfected controls (25,563.82 CTFC ± 6,629.78) (Suppl. Fig. 1 G and H). All PEDF deficient neurons reached the CP by P15, but showed broader distribution of neurons in the CP compared to the control, like that seen following Rpsa KD (Suppl. Fig. 3). The initial analysis revealed that PEDF deficient neurons showed similar morphological defects to Rpsa deficient neurons, with PEDF KD neurons having significantly fewer and shorter extensions with less complex branching as compared to the scramble shRNA control neurons (Fig. 3 and 4A). The mean number of dendrites from the soma in PEDF deficient cells was 2.6 ± 0.21, while the mean number of dendrites from the soma in control neurons was 4.4 ± 0.25 (Fig. 4B). The mean dendrite length in PEDF deficient cells was 27.0 µm ± 2.6 as compared to control neurons with a mean dendrite length of 50.9 µm ± 3.6 (Fig. 4C). These data indicate that PEDF is important for the initiation and elongation of dendrites. Additionally, Sholl analysis of PEDF deficient neurons indicates that these neurons have a statistically significant defect in dendritic branching at radial distances 20-70 µm (Fig. 4D). The Sholl profile is consistent with our analysis of dendrite number and length, since it also indicates that PEDF deficient cells have fewer and shorter dendrites than controls.

**Figure 4.**
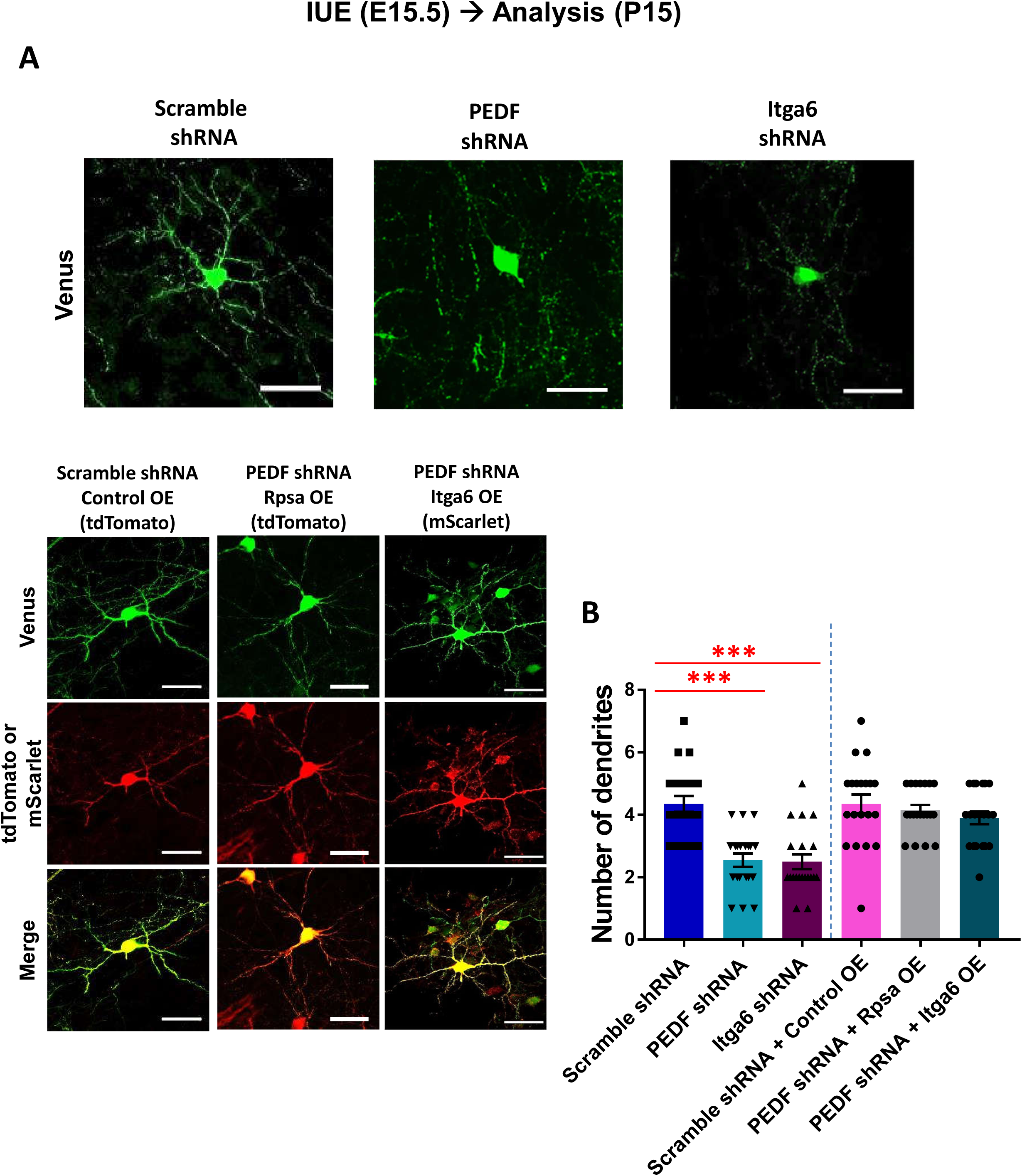
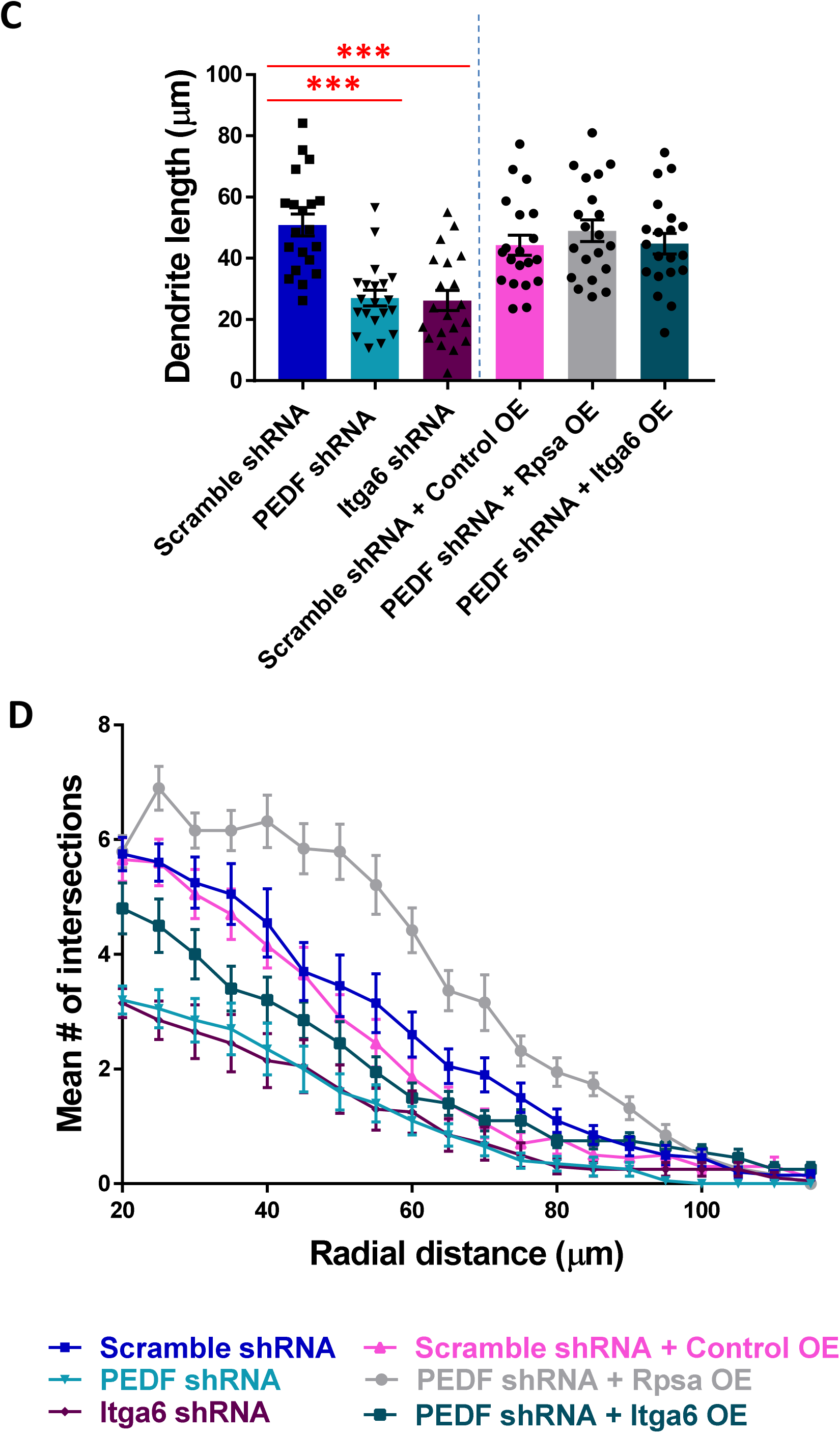
PEDF KD and Itga6 KD result in similar neuromorphological defects and Rpsa OE and Itga6 OE can both rescue defects after PEDF KD. A) Neurons transfected via *in utero* electroporation at E15.5, scale bars = 50 µm. B) One-way ANOVA determined there was a statistically significant difference (F(7,152) = 12.211, *p* < 0.0005). Tukey post hoc test revealed that mean number of dendrites was statistically significantly lower for PEDF shRNA (2.55 ± 0.211, *p*<0.0005) and Itga6 shRNA groups (2.50 ± 0.235, *p*<0.0005) as compared to the scramble shRNA (4.35 ± 0.254). There was no statistically significant difference in mean number of dendrites between the PEDF shRNA + Rpsa OE (4.15 ± 0.167, *p*=0.999) and PEDF shRNA + Itga6 OE (3.90 ± 0.204, *p*=0.874) as compared to the scramble shRNA + control OE group (4.35 ± 0.302). For all groups n=20 cells from 3 mice. Data are represented as mean ± SEM. C) One-way ANOVA determined there was a statistically significant difference (F(7,152) = 16.134, *p* < 0.0005). Tukey post hoc test revealed that mean dendrite length was statistically significantly lower for PEDF shRNA (27.04 ± 2.56, *p*<0.0005) and Itga6 shRNA (26.23 ± 3.30, *p*<0.0005) groups, as compared to the scramble shRNA group (50.86 ± 3.56). There was no statistically significant difference between PEDF shRNA + Rpsa OE (48.99 ± 3.569, *p*=0.956) and PEDF shRNA + Itga6 OE (44.76 ± 3.366, *p*=1.000) as compared to the scramble shRNA + control OE group (44.265 ± 3.307). For all groups n=20 cells from 3 mice. Data are represented as mean ± SEM. D) Sholl analysis shows dramatic decrease in branching in PEDF and Itga6 deficient cells that is rescued in the PEDF shRNA + Rpsa OE and PEDF shRNA + Itga6 OE groups. Two-way ANOVA with Dunnet multiple comparison test shows a significant decrease in branching for the PEDF shRNA and Itga6 shRNA groups at radial distances 20-70 µm as compared to the scramble shRNA group, F(133,3040) = 4.905, *p*<0.0001. For all groups n=20 cells from 3 mice.

To determine whether these similar phenotypes could be a result of PEDF and Rpsa functioning in the same signaling pathway we overexpressed Rpsa in PEDF deficient cells. Rpsa OE rescued morphological defects in the initiation and elongation of dendrites after PEDF KD *in vivo* (Fig. 4A). The mean number of dendrites from the soma in PEDF KD + Rpsa OE neurons was 4.2 ± 0.17, which was not significantly different from the mean of 4.35 ± 0.30 for the double transfected control neurons (scramble shRNA + control OE) (Fig. 4B). The mean dendrite length in PEDF KD + Rpsa OE neurons was 44.3 µm ± 3.6, which was not significantly different from the double transfected control neurons (scramble shRNA + control OE) with a mean dendrite length of 44.3 µm ± 3.3 (Fig. 4C). These data indicate that PEDF is involved in Rpsa-mediated initiation and elongation of dendrites. Additionally, Sholl analysis of PEDF KD + Rpsa OE neurons indicates that Rpsa OE was able to compensate for the defect in branching complexity caused by PEDF KD and resulted in an increase in branching (Fig. 4D). Taken together with the fact that PEDF is a ligand of Rpsa, these results suggest that PEDF binding initiates Rpsa signaling to regulate neuronal morphogenesis.

### Rpsa and Itga6 show similar defects in neuronal morphology after KD and Itga6 OE can rescue morphological defects after PEDF KD

Rpsa is known to bind to Itga6 on the plasma membrane (25). To determine if Itga6 is also involved in PEDF-Rpsa signaling, we used Itga6 shRNA in combination with *in utero* electroporation at E15.5 to conduct an initial analysis of Itga6 deficient neurons *in vivo*. KD efficiency of the Itga6 shRNA was tested via Western Blot and the knockdown efficiency was approximately 79% (Suppl. Fig. 1A and B). In addition, we performed cell staining for Itga6 in primary cortical neurons that were transfected with Itga6 shRNA. Neurons positive for Itga6 shRNA (3,947.66 CTFC ± 819.61) had significantly decreased fluorescence intensity for Itga6, as compared with untransfected controls (21,478.55 CTFC ± 6,285.73) (Suppl. Fig. 1 I and J). Itga6 deficient neurons did not show any deficits in layering with all Itga6 KD neurons reaching the CP by P15 (Suppl. Fig. 3). The analysis revealed that Itga6 deficient neurons showed similar defects to Rpsa and PEDF deficient neurons in neuronal morphogenesis (Fig. 3 and 4A). The mean number of dendrites from the soma in Itga6 deficient cells was 2.5 ± 0.24, while the mean number of dendrites from the soma in control neurons was 4.4 ± 0.25 (Fig. 4B). The mean dendrite length in Itga6 deficient cells was 34.3 µm ± 2.7 as compared to control neurons with a mean dendrite length of 50.9 µm ± 3.6 (Fig. 4C). These data indicate that Itga6 is a key protein for the initiation and elongation of dendrites. Additionally, Sholl analysis of Itga6 deficient neurons indicates that these neurons have a statistically significant defect in dendritic branching at radial distances 20-70 µm, similar to the defects observed in neurons deficient in Rpsa and PEDF (Fig. 3D and 4D).

To determine whether Itga6 is involved in PEDF signaling, we overexpressed Itga6 in PEDF deficient cells. We found that Itga6 OE in PEDF deficient neurons can rescue the morphological defects associated with PEDF KD (Fig. 4A). This rescue phenotype was like the control (scramble shRNA + control OE) and PEDF shRNA + Rpsa OE phenotypes. Thus, these results suggest that Itga6 is involved in a PEDF-Rpsa signaling pathway. The mean number of dendrites from the soma in PEDF KD + Itga6 OE neurons was 4.1 ± 0.29, which was not significantly different from the mean of 4.35 ± 0.30 for the double transfected control neurons (scramble shRNA + control OE) (Fig. 4B). The mean dendrite length in PEDF KD + Itga6 OE neurons was 44.8 µm ± 3.4, which was not significantly different from the double transfected control neurons (scramble shRNA + control OE) with a mean dendrite length of 44.3 µm ± 3.3 (Fig. 4C). This indicates that Itga6 is involved both in the initiation and elongation of dendrites mediated by PEDF and Rpsa. Additionally, Sholl analysis of PEDF KD + Itga6 OE neurons indicates that the branching complexity is restored (Fig. 4D). These results suggest that Itga6, a known binding partner of Rpsa, is important for PEDF-Rpsa signaling to regulate dendrite formation during initiation and elongation stages, and dendritic branching.

### Itga6 OE increases and stabilizes Rpsa expression on the plasma membrane by preventing ubiquitination of Rpsa

Rpsa and Itga6 expression have been shown to be co-regulated and Rpsa and Itga6 co-localize in the cytosol, possibly leading to their being trafficked together to the plasma membrane (24, 25). We tested whether Itga6 could be increasing Rpsa expression on the plasma membrane, which would allow PEDF-Rpsa signaling to occur more efficiently. We transfected 6XHis-Rpsa (only Rpsa OE) and 6XHis-Rpsa + FLAG-Itga6 (both Rpsa and Itga6 OE) into COS-1 cells and performed subcellular fraction to isolate the cytosolic and membrane fractions. Then, Rpsa expression level in the 6XHis-Rpsa + FLAG-Itga6 group was analyzed and compared to the control OE group and Rpsa OE only group. We found that Rpsa expression on the plasma membrane was increased with Itga6 OE, as compared to when only Rpsa was overexpressed (Fig. 5A). This suggests that Itga6 could be contributing to PEDF-Rpsa signaling to regulate neuronal morphology by increasing the amount of Rpsa available on the plasma membrane to participate in signaling and may explain the similar neuronal morphology phenotypes following Rpsa KD, PEDF KD, and Itga6 KD.

**Figure 5.**
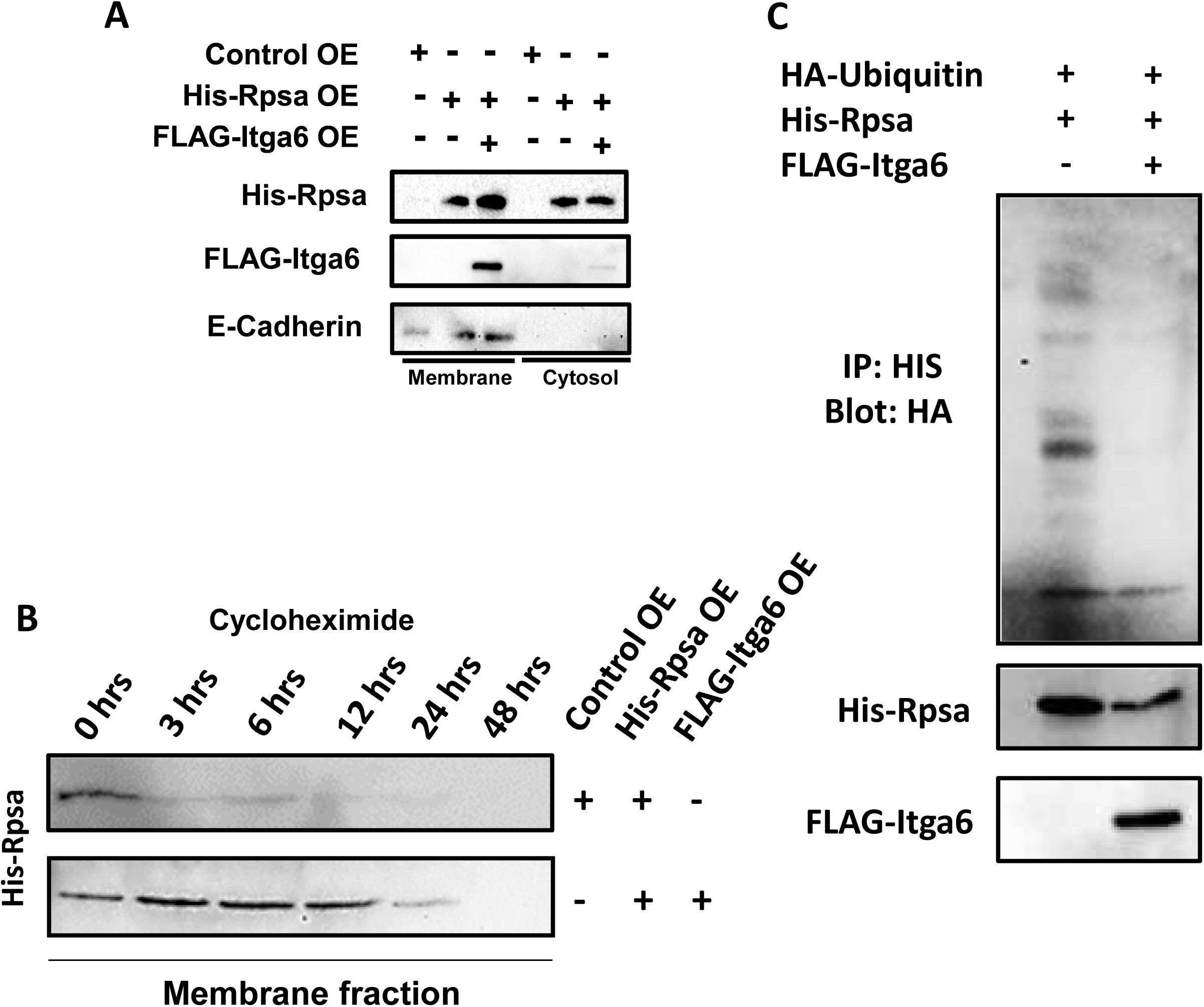
Itga6 OE increases and stabilizes Rpsa expression on the plasma membrane. A) Western blot showing Rpsa and Itga6 expression in the membrane and cytosolic subcellular fractions after either empty backbone control plasmid OE, Rpsa OE, or Rpsa OE + Itga6 OE and sub-cellular fractionation. E-cadherin was used as a membrane fraction marker to show that the sub-cellular fractionation was effective. B) Rpsa expression levels on the plasma membrane of N-2a cells. Sub-cellular fractionation was completed to isolate the membrane fraction. Fractionation was performed at 0, 3, 6, 12, 24, and 48 hours following cycloheximide treatment. C) Pull-down assay showing decreased ubiquitination in Rpsa OE + Itga6 OE cell lysate, as compared to Rpsa OE only lysate.

Since Itga6 OE was shown to increase the amount of Rpsa present in the membrane, we next tested if Itga6 OE could increase the time spent by Rpsa in the membrane before internalization or degradation. We transfected Neuro2a (N-2a) mouse neuroblastoma cells with either 6XHis-Rpsa (only Rpsa OE) or 6XHis-Rpsa + FLAG-Itga6 (both Rpsa and Itga6 OE) and 48 hours after transfection treated the cells with cycloheximide, which inhibits protein synthesis. Cells were then subject to sub-cellular fractionation to isolate the membrane fraction at 0, 3, 6, 12, 24, and 48 hours following cycloheximide treatment. Western blot for 6XHis-Rpsa using anti-His antibody was completed on the membrane fraction revealing that Rpsa remained present in the membrane for dramatically longer following cycloheximide treatment when both Rpsa and Itga6 were overexpressed, as compared with when only Rpsa was overexpressed (Fig. 5B). This suggests that Itga6 stabilizes Rpsa in the membrane.

To determine the mechanism by which Itga6 increases and stabilizes Rpsa expression on the plasma membrane, we transfected N-2a cells with HA-ubiquitin and either 6XHis-Rpsa (only Rpsa OE) or 6XHis-Rpsa + FLAG-Itga6 (both Rpsa and Itga6 OE). Ubiquitin has already been shown to regulate the presence of Rpsa at the plasma membrane (31). Pull-down assay was done 48 hours after transfection to isolate 6XHis-Rpsa. Western blot using anti-HA antibody showed decreased ubiquitination when both Itga6 and Rpsa were overexpressed, as compared to when only Rpsa was overexpressed (Fig. 5C). This suggests that Itga6 stabilizes the expression of Rpsa on the plasma membrane by preventing its ubiquitination, which would lead to internalization or degradation.

### Rpsa KD and PEDF KD, but not Itga6 KD, cause a decrease in overall spine density and a change in spine morphology

We further investigated the neuromorphological changes observed after Rpsa KD, PEDF KD, and Itga6 KD by determining if there was a change in dendritic spine density and morphology, since these factors can impact synaptic function. We used *in utero* electroporation at E15.5 to transfect the relevant plasmids and analyzed spines at P15 (Fig. 6A). We observed a decrease in overall spine density after Rpsa KD (0.28 ± 0.04) and PEDF KD (0.21 ±. 04), as compared to the control CRISPR (0.70 ± 0.05) and scramble shRNA (0.97 ± 0.09) controls, respectively (Fig. 6B). The Rpsa KD and PEDF KD groups also showed significant differences in spine morphology (Fig. 6C). There was a significant decrease in mushroom spines following both Rpsa KD (5.88%) and PEDF KD (6.98%) as compared to control CRISPR (55.56%) and scramble shRNA (60.10%), respectively. Thin spines were significantly increased after Rpsa KD (66.18%), while thin spines were significantly decreased following PEDF KD (34.88%) as compared to control CRISPR (22.22%) and scramble shRNA (20.21%), respectively. Rpsa KD also resulted in a significant decrease in stubby spines (10.29%), as compared to control CRISPR (18.06%), and a significant increase in filopodia spines (16.18%) as compared to control CRISPR (2.78%). This indicates that Rpsa KD results in a clear shift towards a more immature spine morphology, while PEDF KD follows this same trend. Interestingly, Itga6 KD (1.01 ± 0.10) resulted in a spine density phenotype like that of the scramble shRNA control group (Fig. 6B). The Itga6 KD and scramble shRNA groups also showed similar spine morphology phenotypes, with no significant differences in spine morphology between these groups (Fig. 6C). This suggests that PEDF-Rpsa signaling regulates spine density in addition to morphology, while Itga6 may not be involved in the regulation of spine formation by PEDF-Rpsa signaling.

**Figure 6.**
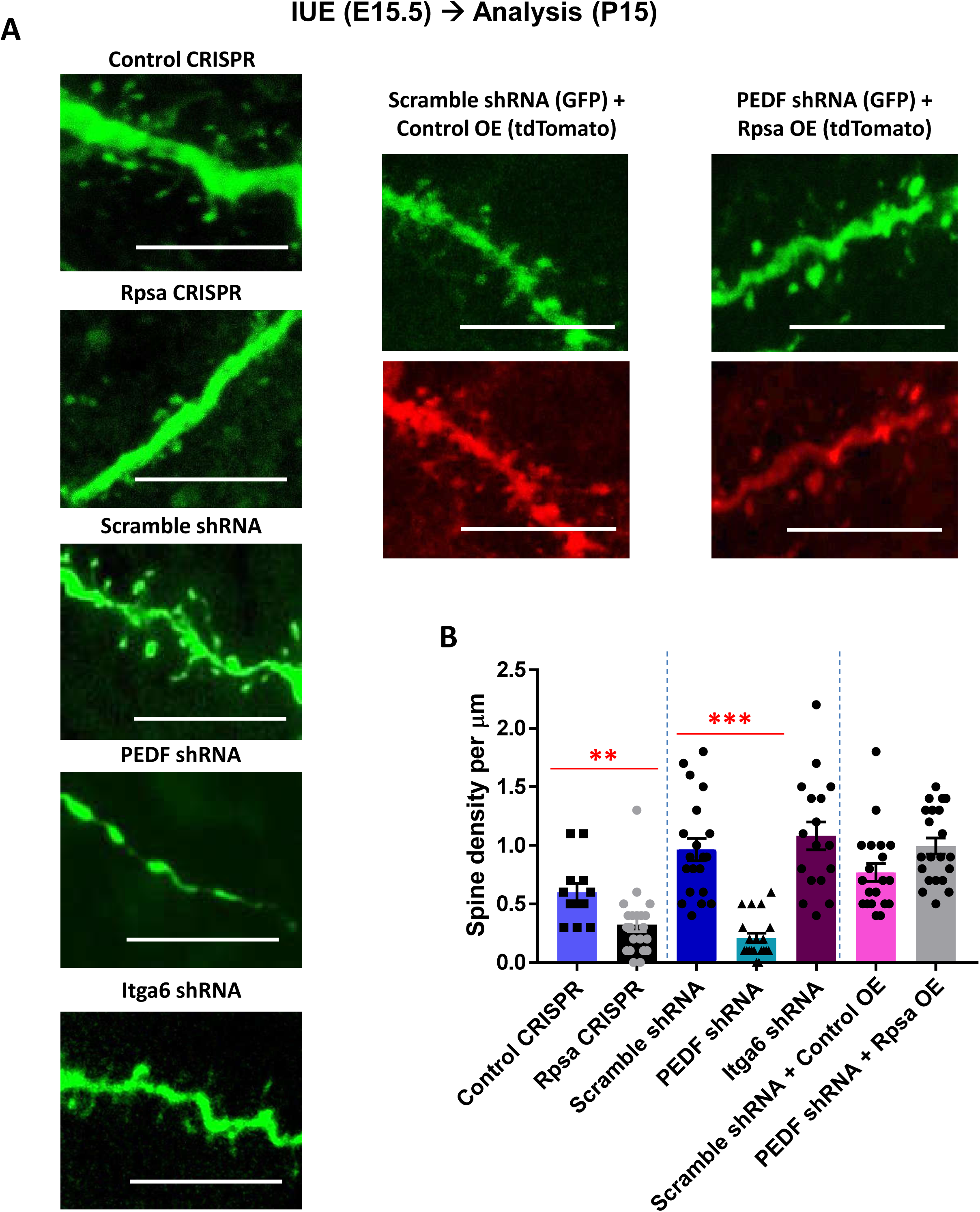
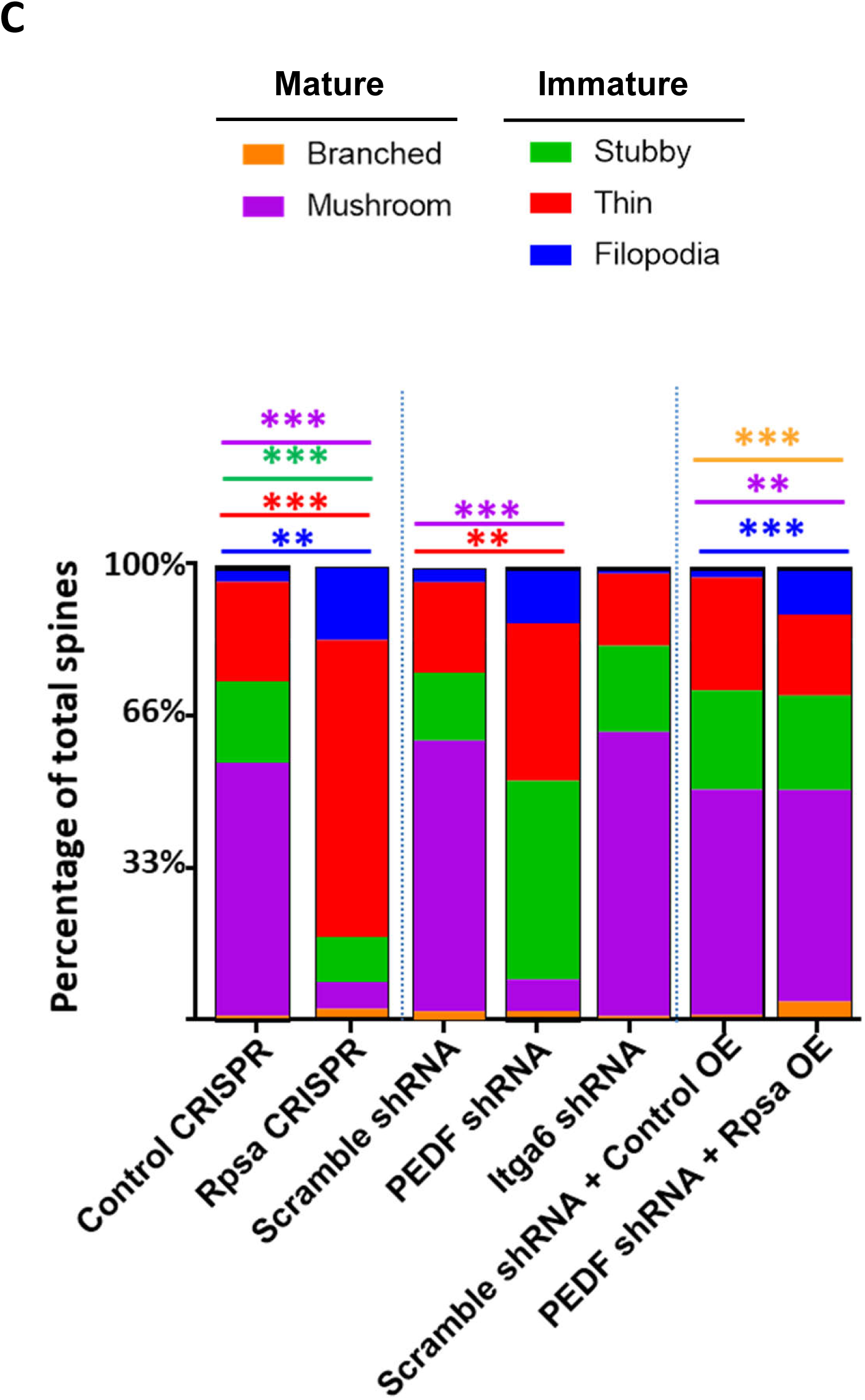
Rpsa KD and PEDF KD, but not Itga6 KD, cause a decrease in overall spine density and a change in spine morphology. A) Neurons transfected via *in utero* electroporation at E15.5, images taken with 63x objective and 5x zoom to visualize spines, scale bares = 5 µm. B) One-way ANOVA determined there was a statistically significant difference (F(6,127) = 23.321, *p* < 0.0005). Tukey post hoc test revealed that mean spine density per µm was statistically significantly lower for Rpsa CRISPR (0.28 ± 0.04, n=21) as compared to control CRISPR (0.70 ± 0.05, n=12) (*p*=0.001) and mean spine density per µm was statistically significantly lower for PEDF shRNA (0.21 ± 0.04, n=20) as compared to scramble shRNA (0.97 ± 0.09, n=20) (*p*<0.0005). However, the Itga6 shRNA group (1.01 ± 0.10, n=17) did not show a significant difference compared to the scramble shRNA group (*p*=0.999). The PEDF shRNA + Rpsa OE group (0.99 ± 0.07, n=21) was not significantly different from the scramble shRNA + control OE group (0.77 ± 0.08, n=20) (*p*=0.230). Data are represented as mean ± SEM. For each group, spines were analyzed from 3 mice. C) Graph showing differences in percentage of total spines with filopodia, thin, stubby, mushroom, or branched morphology for each experimental group. Statistical analyses were done using raw spine density per µm data. Branched spines were significantly increased in the PEDF shRNA + Rpsa OE group (0.0381 ± 0.01, n=21), as compared to scramble shRNA + control OE (0.0050 ± 0.01, n=20), t(408) = −2.295, *p*<0.0005. Mushroom spines were significantly decreased in the Rpsa CRISPR (0.0190 ± 0.01, n=21, *p*<0.0005) and PEDF shRNA (0.0150 ± 0.01, n=20, *p*<0.0005) groups as compared to control CRISPR (0.3333 ± 0.04, n=12) and scramble shRNA (0.5800 ± 0.03, n=20), respectively, while mushroom spines were significantly increased in the PEDF shRNA + Rpsa OE (0.4619 ± 0.03, n=21, *p*=0.004) group compared to the scramble shRNA + control OE (0.3850 ± 0.03, n=20). Stubby spines were significantly decreased in the Rpsa CRISPR (0.0333 ± 0.01, n=21, *p*<0.0005) compared to control CRISPR (0.1083 ± 0.03, n=12), t(328) = 2.770. Thin spines were significantly increased in the Rpsa CRISPR group (0.2143 ± 0.03, n=21) as compared to control CRISPR (0.1333 ± 0.03, n=12) (t(328) = −1.826, *p*<0.0005), while thin spines were significantly decreased in the PEDF shRNA group (0.0750 ± 0.02, n=20, *p*=0.002), as compared to scramble shRNA (0.1950 ± 0.03, n=20) (F(2,567) = 6.507, *p*=0.002). Filopodia spines were significantly increased in the Rpsa CRISPR (0.0524 ± 0.02, n=21, *p*=0.001) and PEDF shRNA + Rpsa OE (0.1000 ± 0.02, n=21) groups, as compared to the control CRISPR (0.0167 ± 0.01, n=12, *p*<0.0005) and scramble shRNA + control OE (0.0150 ± 0.01, n=20), respectively.

To determine whether Rpsa OE could rescue spine defects after PEDF KD, we used *in utero* electroporation at E15.5 to overexpress Rpsa in PEDF deficient cells (Fig. 6A). The PEDF shRNA + Rpsa OE group (0.99 ± 0.07) showed an overall spine density that was comparable to the scramble shRNA + control OE group (0.77 ± 0.08) (Fig. 6B). The PEDF shRNA + Rpsa OE group showed a significant increase in branched (3.88%), mushroom (47.09%), and filopodia (10.19%) spines compared to the scramble shRNA + control OE group (branched: 0.65%, mushroom: 50.00%, filopodia: 1.95%) (Fig. 6C). This rescue shows an increase in branched and mushroom spines, which indicate a mature morphology, while also showing an increase in filopodia spines, which are considered immature. Thus, Rpsa OE can rescue spine density after PEDF KD, and it is suggested that Rpsa could accelerate the creation of new spines.

### Calcium imaging of brain slices shows sub-threshold functional difference in Rpsa KD neurons

We next investigated whether the Rpsa KD associated decrease in dendrite number and length, as well as reduced dendritic arborization and change in spine density and morphology could be linked to functional deficits, since neuronal morphology is a critical factor in determining the extent of inputs received by the neuron. To investigate function in Rpsa deficient neurons we conducted calcium imaging using live brain slices after *in utero* electroporation at E15.5 collected from male and female mice (approximately P30). The calcium indicator GCaMP6s was used to image neurons double positive for Rpsa or control CRISPR (tdTomato) and GCaMP6s (GFP). GCaMP6s has a relatively high sensitivity and is a well-validated calcium indicator (32). Rpsa deficient neurons had a significantly lower mean difference between maximum and minimum peaks (6.7 ±1.8), as compared with control neurons (12.8 ± 1.1) when spontaneous calcium activity was measured (Fig. 7A), indicating that the Rpsa deficient neurons show significantly decreased calcium signaling. We found that Rpsa KD neurons showed almost no fluctuation in fluorescence intensity, while the control CRISPR neurons showed a dramatically greater change in fluorescence intensity (Fig. 7B). Since few action potentials were recorded, these differences in fluorescence intensity indicate that Rpsa KD neurons have a defect in calcium signaling at a sub-threshold level. Calcium signaling directly relates to neuronal function by contributing to transmission of the depolarizing signal, synaptic signaling processes, neuronal energy metabolism, and neurotransmission (33).

**Figure 7.**
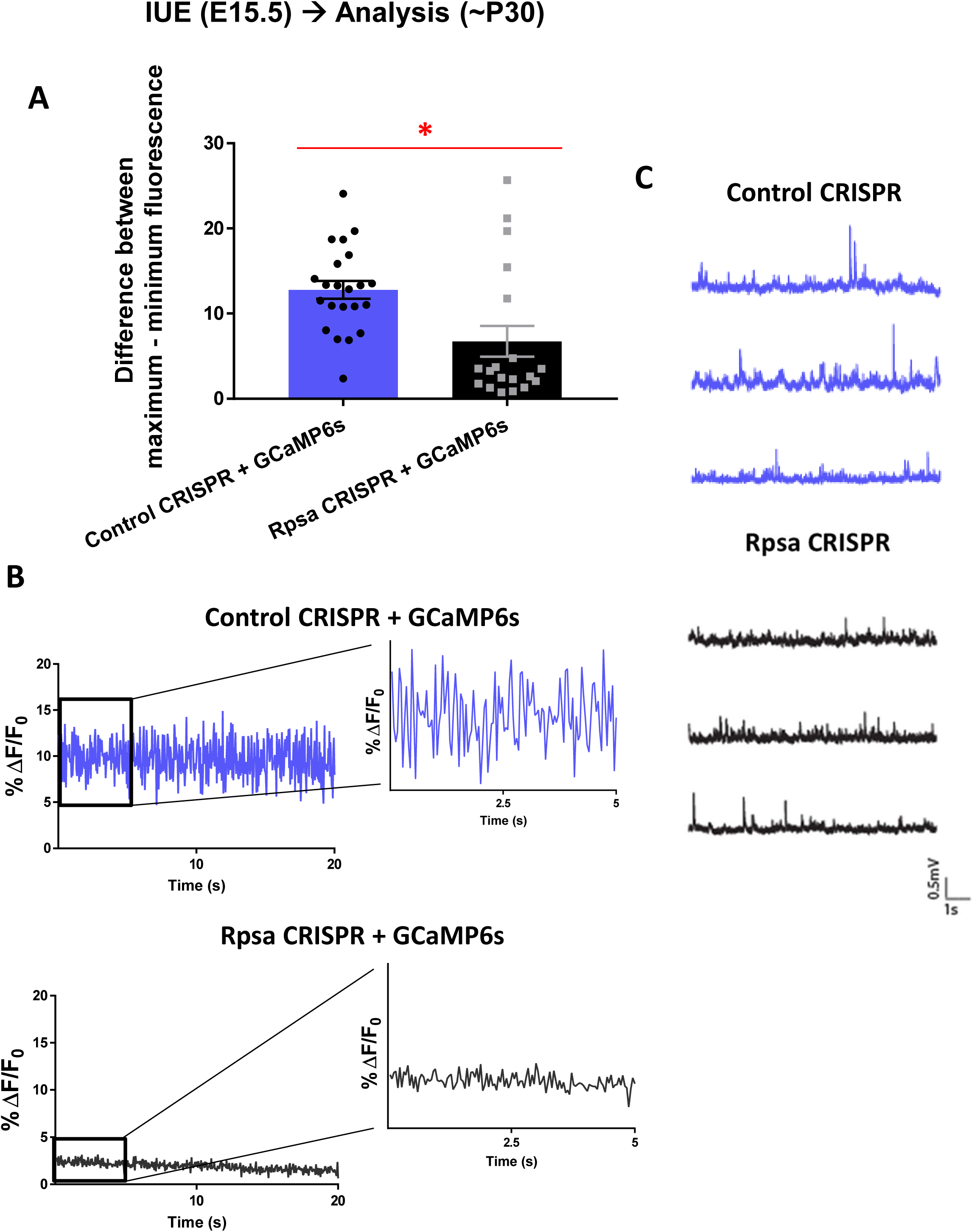
Calcium imaging shows sub-threshold functional difference in Rpsa KD neurons. A) Mean difference in GCaMP6s fluorescence intensity. Two-tailed independent-samples t-test showed the average difference between maximum and minimum peaks was significantly greater in the control (12.79 ± 1.06, n=22 cells from 3 mice) compared to Rpsa KD (6.73 ± 1.80, n=19 cells from 3 mice), t(39) = 2.992, *p* = 0.035. Data are represented as mean ± SEM. B) Representative % change in fluorescence (ΔF/F_0_) for representative individual neuron in mouse brain slice. Single action potentials are defined as ΔF/F0 ≥ 23% ± 3.2%. The left panels were taken over 20 seconds and the right panels show a 5 second section of each representative trace, with the y-axis measuring 8 units to provide a zoomed in view. C) Electrophysiology recordings confirm low spontaneous firing rate of control CRISPR and Rpsa CRISPR neurons. Representative traces of spontaneous activity of 3 control CRISPR and Rpsa CRISPR neurons at resting membrane potential in current-clamp mode. n= 9 cells from 3 control CRISPR mice and n= 7 cells from 2 Rpsa CRISPR mice.

To complement our calcium imaging data, we conducted whole-cell electrophysiological recordings in mice (approximately P30) after knocking down Rpsa by CRISPR by *in utero* electroporation at E15.5. In agreement with our calcium imaging results, electrophysiological recordings showed that control and Rpsa KD neurons do not fire action potentials spontaneously at rest (Fig. 7C). However, both control and Rpsa deficient neurons fired action potentials upon stimulation (Suppl. Fig. 4). Intrinsic membrane properties were comparable between groups and only differed significantly in capacitance, indicating a difference in cell size (Suppl. Fig. 4 and Table 1). Taken together, this corroborates our calcium imaging results by confirming the low spontaneous firing rate in both control and Rpsa KD cells and showing that Rpsa KD neurons are healthy and able to fire action potentials.

**Table 1.** Comparison of membrane properties between Control CRISPR and Rpsa CRISPR

|  |  |  |  |  |  |  |  |  |
| --- | --- | --- | --- | --- | --- | --- | --- | --- |
|  | V<sub>m</sub><br>(mV) | Resistance<br>(MΩ) | τ (ms) | C (pF) | Rheobase<br>(pA) | Spike<br>height<br>(mV) | Overshoot<br>(mV) | Spike<br>threshold<br>(mV) |
| Control<br>CRISPR<br>(n=9) | -67 ±<br>1.2 | 60 ± 4.9 | 14 ±<br>1.2 | 251 ±<br>30.2 | 141 ±<br>10.5 | 84 ±<br>1.8 | 50 ± 1.2 | -34 ± 1.1 |
| Rpsa<br>CRISPR<br>(n=7) | -64 ±<br>0.9 | 50 ± 5.1 | 18 ±<br>2.6 | 369 ±<br>35.8 | 149 ±<br>33.3 | 81 ±<br>1.7 | 48 ± 2.4 | -32 ± 1.2 |

V<sub>m</sub> = membrane potential, τ = membrane time constant, C = capacitance
Data are represented as mean ± SEM.

## Discussion

Our study reports a role for Rpsa in regulating neuronal morphogenesis, including apical dendrite orientation, dendrite initiation and extension, dendritic branching, and spine density and morphology and suggests that morphological defects caused by Rpsa KD have sub-threshold functional consequences for calcium signaling. Additionally, we report that Itga6 increases and stabilizes Rpsa expression on the plasma membrane by preventing the ubiquitination of Rpsa. By performing *in vivo* rescue experiments, we found that PEDF-Rpsa-Itga6 signaling is essential for neuromorphogenesis.

Orientation of the apical dendrite and neuronal morphogenesis was dramatically impacted by disrupting Rpsa signaling. Improper orientation of the apical dendrite may have led to formation of inappropriate synaptic connections, since the apical dendrite may not have been within sufficient vicinity of its target neuron to allow synaptogenesis to occur. The decrease in number of dendrites and dendrite length suggests a deficit in the initiation and elongation of dendrites, respectively. Defects in these early stages of morphogenesis may have caused further defects at later stages such as dendritic arborization, as a severe defect in dendritic branching was observed in Rpsa deficient neurons. Rpsa deficient neurons show aberrant orientation of the apical dendrite at the proximal region of the apical dendrites (Fig. 2). However, the distal region of apical dendrites turns into the proper direction to the marginal zone (Fig. 2A). This implies that Rpsa affects the very initial step of dendrite extension after its initiation from soma, but not the later dendrite extension step. Future studies including time-lapse live imaging will help understand the precise function of Rpsa in regulating dendrite behavior during the initial step of dendrite extension and orientation.

While a significant decrease in dendrite length was seen at P15 (Fig. 3), no difference was observed between Rpsa KD and control CRISPR in dendrite length at P3 (Suppl. Fig. 2). The mean length measurement at P3 is heavily influenced by the length of the apical dendrite since it is one of the only extensions present at this developmental time-point and basal dendrites were not yet formed. Therefore, the defects in dendrite length seen at P15 resulted from the defects in basal dendrites, indicating the importance of Rpsa in basal dendrite formation.

Genetically manipulating the proposed Rpsa signaling pathway using various rescue experiments allowed us to provide convincing evidence that PEDF is the ligand responsible for initiating Rpsa signaling to regulate neuronal morphogenesis and that a Rpsa-Itga6 signaling complex is important for dendrite formation (Fig. 8A). Furthermore, we show that Itga6 increases Rpsa’s expression and stabilization on the plasma membrane, where it must be located to interact with its extracellular ligand PEDF, indicating that Rpsa’s role as a receptor is critical for regulating neuromorphogenesis. Importantly, it has been shown that rat cerebral cortical neurons secrete PEDF, suggesting mouse cortical neurons secrete PEDF since cell staining indicated PEDF expression in mouse cortical neurons (Suppl. Fig. 5A) (34). The morphological defects seen in the PEDF deficient neurons in our *in utero* electroporation experiments suggest that autocrine PEDF-Rpsa signaling is a main pathway in neurons in the developing cortex. Because PEDF is a secreted protein, PEDF can also bind Rpsa receptors present on other neurons to initiate a paracrine signaling pathway, but this paracrine signaling pathway would be less efficient than the autocrine pathway. In our rescue experiment with Rpsa overexpression in PEDF deficient cells, almost all autocrine pathways would be prevented by PEDF KD, but increased Rpsa on the membrane should increase the efficiency of the paracrine signaling pathway by accelerating the opportunity for PEDF to bind to an Rpsa receptor and compensate for the PEDF KD.

**Figure 8.**
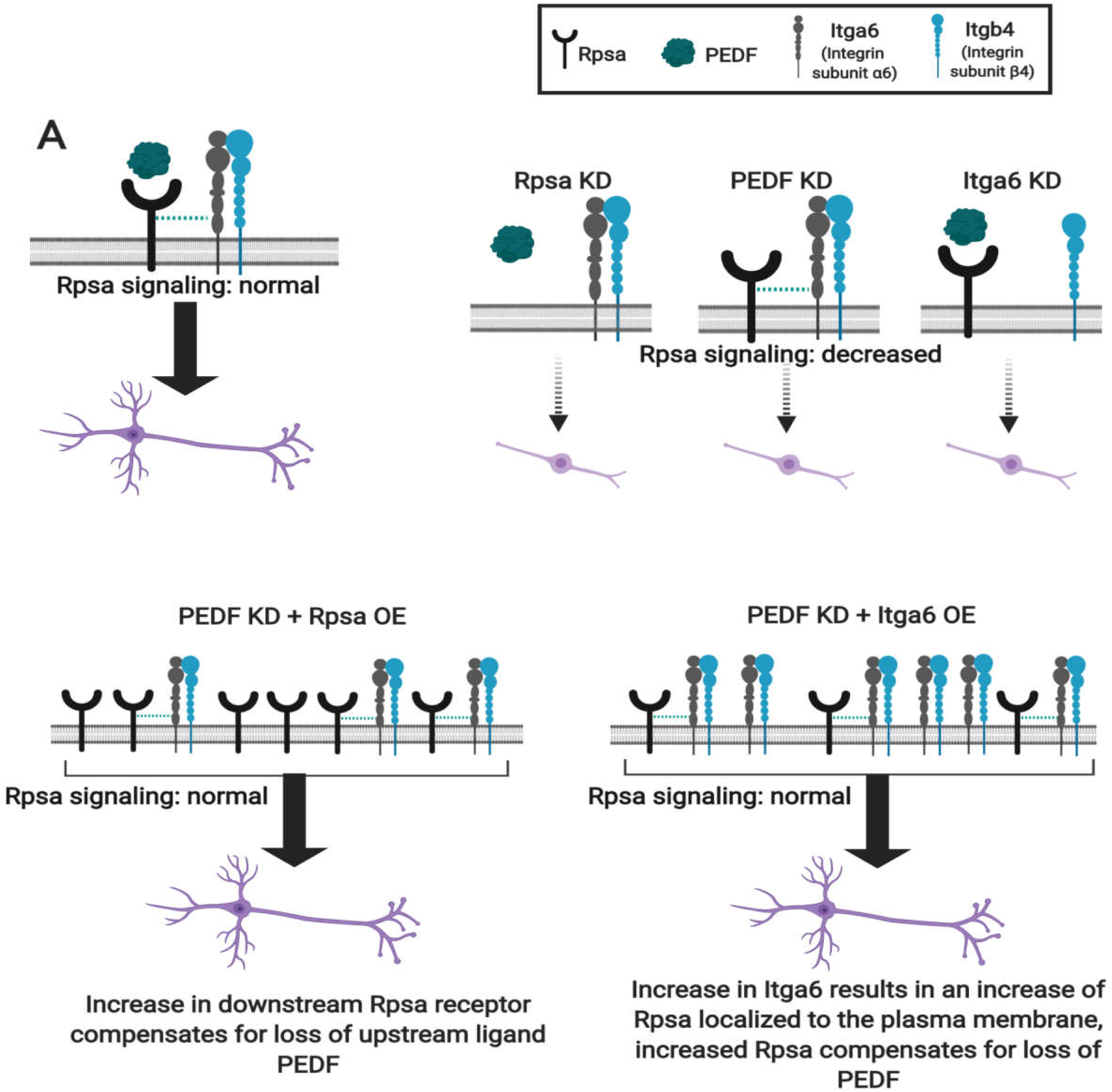
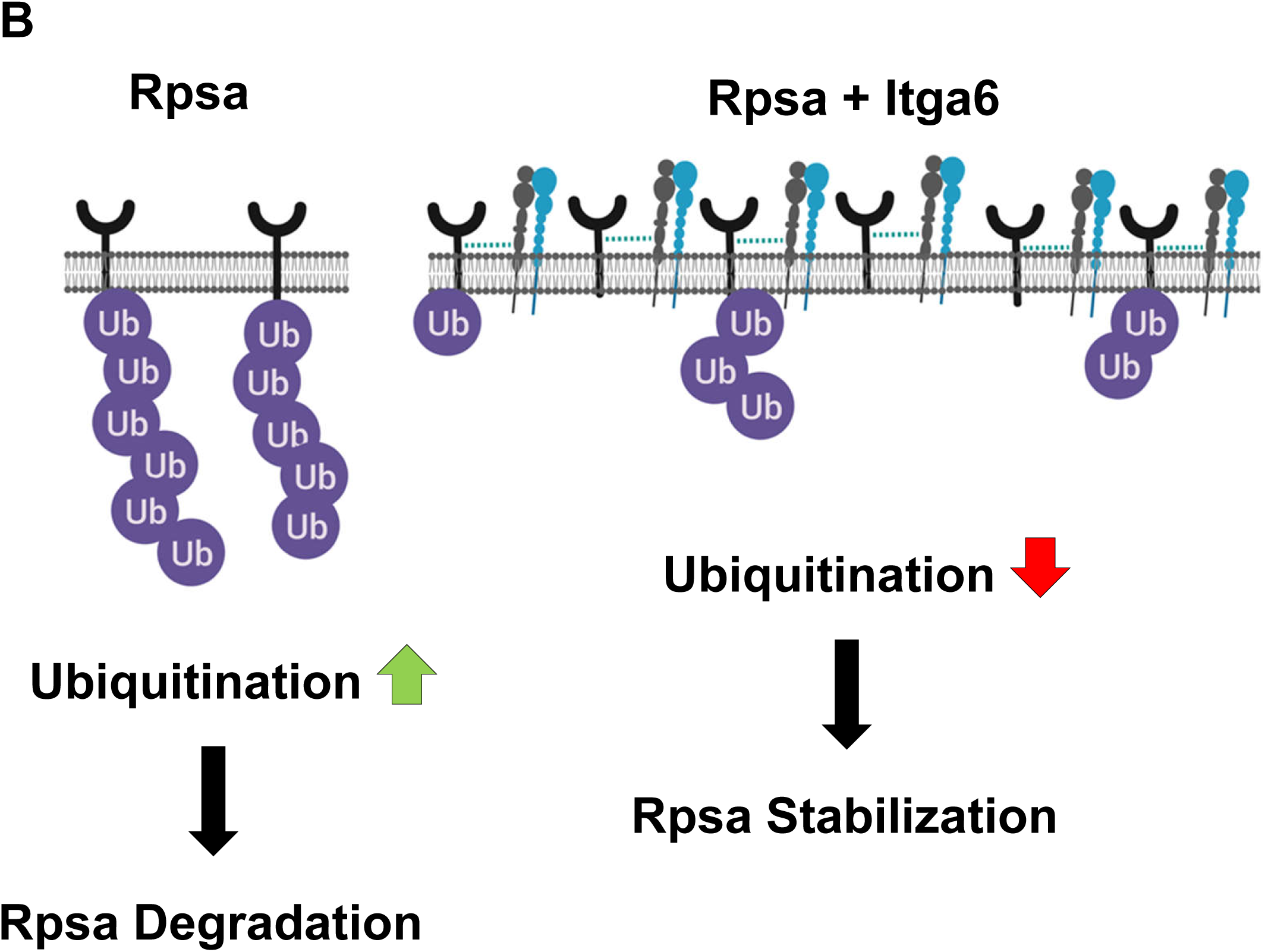
Schematic of Rpsa/PEDF/Itga6 signaling regulating neuronal morphogenesis. A) Under normal conditions Itga6, which is likely bound to a β4 subunit to function as part of a complete integrin, is co-localized on the plasma membrane with Rpsa. Both Rpsa and Itga6 facilitate the binding of PEDF to Rpsa, which results in signaling to regulate proper neuronal morphogenesis (top left). Following Rpsa KD, PEDF KD, or Itga6 KD the signaling mechanism initiated by Rpsa is attenuated, resulting in severe defects in morphology (top right). Knockdown of the upstream ligand PEDF can be compensated for by increasing its downstream receptor Rpsa, to cause increase of Rpsa-Itga6-Itgb4 complex, thus resulting in normal level of signal intensity and normal morphogenesis. The increase in Rpsa-Itga6-Itgb4 complex can also be accomplished by Itga6 OE (bottom). B) Itga6 increases both the expression level and time spent by Rpsa in the plasma membrane by preventing ubiquitination of Rpsa.

Our rescue experiments showed that dendrite formation when Itga6 was overexpressed was similar to dendrite formation when Rpsa was overexpressed. These similar rescue phenotypes agree with our data indicating that an increase in Itga6 expression leads to an increase in Rpsa localization and stabilization on the plasma membrane, producing a similar effect to the Rpsa OE rescue (Fig. 8A). The overexpression of Rpsa and Itga6 in combination with cellular fractionation revealed that Itga6 OE facilitates Rpsa expression on the plasma membrane (Fig. 5A). Itga6 increases both the expression level and time spent by Rpsa in the plasma membrane by preventing ubiquitination of Rpsa (Fig. 8B). Rpsa has a total of 6 ubiquitination sites: Lys11, Lys40, Lys52, Lys57, Lys89, and Lys166. While Lys89 may be located in a computer predicted transmembrane domain at residues 86-101 and Lys166 is likely to be included in the extracellular laminin binding domain, Lys11, Lys40, Lys52, and Lys57 are located inside the cell and are likely targets for ubiquitination (35). Itga6 could potentially bind Rpsa such that Itga6 blocks the ubiquitination site on Rpsa that regulates its degradation. Itga6 is expressed in some of the same regions of the cortex as Rpsa, including the CP (Suppl. Fig. 5B). Previous studies indicate that ubiquitin E3 ligase Nedd4 is primarily responsible for labeling Rpsa for ubiquitin-mediated internalization (36). Thus, Nedd4 is likely involved in PEDF-Rpsa-Itga6 signaling, and the mechanisms of Rpsa ubiquitination by Nedd4 should be analyzed in the future.

The Sholl profiles for both the PEDF shRNA + Rpsa OE and PEDF shRNA + Itga6 OE rescue experiments show an increase in complexity of dendritic branching as compared to the PEDF shRNA group, indicating that overexpression of either Rpsa or Itga6 rescues this aspect of the PEDF KD phenotype (Fig. 4D). However, the PEDF shRNA + Rpsa OE group showed even more complex branching than the scramble shRNA + control OE group. Rpsa OE could have resulted in a more complex branching phenotype, since Rpsa is further downstream than PEDF in the proposed signaling pathway. Thus, overexpression at a more downstream level in the signaling pathway could have overstimulated the mechanism responsible for regulating branching, leading to an increase in branching complexity.

To determine whether the dramatic morphological defects caused by Rpsa KD impact function, we conducted calcium imaging on live brain slices and detected a difference between Rpsa KD cells and control cells in sub-threshold calcium signaling. The GCaMP6s calcium indicator was used, which has a relatively high sensitivity and has commonly been used to image relatively low cytoplasmic calcium levels (32). The fluorescence intensity has a greater percent change for the control, while Rpsa deficient cells maintain a more constant level of fluorescence intensity. Sub-threshold changes in calcium signaling in Rpsa KD neurons could be due to differences in NMDA channel, NMDA receptor (NMDAR), or voltage-gated calcium channels (VGCCs) functions, which could be reduced simply because of the decrease in dendritic surface area by less dendrites and shorter dendrite length (37–41). Many previous studies differ from ours in that they focus on calcium signaling in active spines. However, the amount of calcium that enters via these synaptic mechanisms may be very small, requiring that the signal be greatly amplified by intracellular calcium release to be detected (42). We would be able to detect sub-threshold calcium signaling changes resulting from calcium release from internal stores in the soma. The morphological defects caused by Rpsa KD may be impacting the NMDA, NMDAR, and/or VGCCs responsible for mediating calcium influx, thus resulting in sub-threshold calcium signaling changes.

Single action potentials are defined as ΔF/F_0_ ≥ 23% ± 3.2%. Thus, few action potentials were recorded by our calcium imaging. However, this is not surprising, since pyramidal cells can have a low spontaneous firing rate (43). Our electrophysiology recordings confirm the minimal percent change in fluorescence detected by our calcium imaging by showing that inputs do not summate to a level that would cause action potentials for either the Rpsa KD or control group. This indicates that the specific cortical pyramidal neurons that were transfected via *in utero* electroporation have a low spontaneous firing rate. Thus, we should not expect to see many action potentials recorded in our calcium imaging experiments. Taken together, these data suggest that Rpsa KD causes a functional difference in calcium signaling at sub-threshold levels in cortical pyramidal neurons with a low spontaneous firing rate.

Additionally, our electrophysiology recordings indicate that control CRISPR neurons and Rpsa CRISPR neurons display similar intrinsic membrane properties (Suppl. Fig. 3 and Table 1). This is important to note because the 37-kDa Rpsa precursor is a component of the 40S ribosome and is involved in pre-RNA processing, thus it could be suggested that the morphological defects in Rpsa deficient cells may be attributed to a poor overall state of the neurons (5, 44). However, our electrophysiology results suggest that Rpsa deficient cells remain healthy, despite the phenotypic changes associated with Rpsa KD (Suppl. Fig. 3 and Table 1).

These results are of clinical interest because the *Serpinf1* gene, coding for PEDF protein, is in the chromosome 17p13.3 region, which is often deleted causing Miller-Dieker Syndrome (MDS) and duplicated causing the 17p13.3 microduplication syndrome. There is no doubt that neuronal migration defects are a main cause of MDS, but the pathogenesis of MDS has not been completely clarified in detail. MDS has a complex etiology because it caused by a microdeletion that could include more than 26 genes, including *Serpinf1*. While most of the genes deleted in MDS patients have not been investigated, previous studies have analyzed some genes, including *Pafah1b1* (Lis1), *Ywhae* (14-3-3ε) and *Crk* in the MDS critical region (22, 45-49). However, these studies focused on the gene functions in neuronal migration and neural activity, since most MDS patients suffer from epilepsy. The role of neuronal morphogenesis in MDS etiology remains uncertain. PEDF deficiency may not be the main cause of MDS, but it could contribute to MDS pathogenesis. Thus, our findings will provide new insights into the mechanisms underlying MDS.

In conclusion, Rpsa signaling mediates functionally relevant aspects of cortical neuronal morphogenesis including apical dendrite orientation, initiation and elongation of dendrites, dendritic branching, and dendritic spine density and morphology. This Rpsa signaling mechanism is initiated by binding of its ligand PEDF, while Itga6 promotes and stabilizes the expression of Rpsa on the membrane. Future studies should focus on investigating the mechanism by which Rpsa KD impacts the synapse and elucidating the downstream targets of Rpsa during cortical neuronal morphogenesis. The PI3K and MAPK signaling pathways have been shown to be downstream of Rpsa and Itga6 in promoting pancreatic cancer invasion and metastasis (50). This indicates that PI3K and MAPK are potentially downstream of PEDF-Rpsa-Itga6 signaling during cortical development. Furthermore, Nedd4, which is known to ubiquitinate Rpsa, has been implicated in branching and regulation of neurite growth by acting as a downstream target of PI3K/PTEN-mTORC1 (36, 51, 52). Future studies should determine if these signaling mechanisms are downstream effectors of PEDF-Rpsa-Itga6 mediated regulation of cortical neuromorphogenesis. Further investigation of this pathway will advance our understanding of neuronal morphogenesis in normal brain development as well as increase our knowledge of neurodevelopmental disorders, such as MDS.

## Materials and Methods

### Mice

All animal experiments were performed under protocols approved by the Drexel University Animal Care and Use Committees and following the guidelines provided by the US National Institutes of Health. ICR mice were purchased from Taconic Inc. Embryonic day (E) 0.5 was defined as noon of the day the vaginal plug appeared. Females and males were used for *in utero* electroporation and primary cortical neuron culture.

### Plasmids

pCAG-eCas9-GFP-U6-gRNA was a gift from Jizhong Zou (Addgene plasmid # 79145; http://n2t.net/addgene:79145; RRID:Addgene_79145). This plasmid contains high-fidelity eSpCas9 to reduce the off-target effects. gRNAs were designed using the web-based design tools, including CHOPCHOP (http://chopchop.cbu.uib.no/) and CRISPy-web (https://crispy.secondarymetabolites.org/#/input), and then cloned into pCAG-eCas9-GFP-U6-gRNA. Rpsa target gRNA sequence is ATCTACAAAAGGAAAAGTGA(CGG) (112-131 of mouse Rpsa). CRISPR was chosen as the method to accomplish Rpsa KD, as opposed to shRNA, because we were not able to identify a specific shRNA target sequence using web-based design tools (InvivoGen siRNA Wizard, and Invitrogen Block-iT RNAi Designer). For calcium imaging, GFP in pCAG-eCas9-GFP-U6-control-gRNA and pCAG-eCas9-GFP-U6-Rpsa-gRNA was replaced into tdTomato by PCR. 6XHis-tagged Rpsa was cloned into pCAGItdTomato vector (pCAGItdTomato-Rpsa) as described previously (53). CRISPR-resistant Rpsa overexpression plasmid was created by PCR. The sense and antisense primers containing the CRISPR target sequence with mutations were designed, and PCR was performed using PrimeSTAR GXL (Takara) with pCAGItdTomato-Rpsa. Ten mutations without amino acid change were introduced into primers. The sequence with mutation is AT<u>T</u>TA<u>T</u>AA<u>GC</u>G<u>C</u>AA<u>GTCA</u>GA(TGG) where underlined nucleotides are mutated. The insertion of mutations was confirmed by sequencing. To create pCAGEN-GCaMP6s, pCAGEN and pGP-CMV-GCaMP6s were obtained from Addgene, and GCaMP6s fragment amplified by PCR was cloned into pCAGEN. pGP-CMV-GCaMP6s was a gift from Douglas Kim (Addgene plasmid # 40753; http://n2t.net/addgene:40753; RRID:Addgene_40753)(32). pCAGEN was a gift from Connie Cepko (Addgene plasmid # 11160; http://n2t.net/addgene:11160; RRID:Addgene_11160) (54). PEDF and Itga6 shRNAs were designed using the web-based tools described above and cloned into pSCV2-Venus plasmid as described previously (53, 55). The target sequence of mouse PEDF is GAACTTGACCATGATAGAA (849-867). The target sequence for mouse Itga6 shRNA is GACCAAAGACTCGATGTTT (1113-1131). Certified scramble shRNA (ACTACCGTTGTTATAGGTG) was also used as a negative control (Invitrogen). All plasmids used in this study were purified by NucleoBond Xtra purification kit (MACHEREY-NAGEL). pLenti-CMV-Itga6-Myc-DDK-P2A-Puro vector was purchased from Origene Technologies, Inc. (PS100092). Itga6-Myc-DDk fragment was amplified by PCR and cloned into pLV-CAG-P2A-mScarlet plasmid. HA-Ubiquitin was a gift from Edward Yeh (Addgene plasmid # 18712; http://n2t.net/addgene:18712; RRID:Addgene_18712) (56).

### Analysis of genomic alterations by CRISPR/Cas9

pCAG-eCas9-GFP-U6-Rpsa-gRNA plasmid was transfected into mouse Neuro-2a cells using PolyJet transfection reagent (SignaGen Laboratories), and Genomic DNA was isolated and subjected to PCR to amplify the 433 bp fragment containing gRNA target sequence using Q5 High Fidelity DNA polymerase (NEB) and primers (GAATTC(EcoRI)/GAGTTCTAGTGTCAGAAGAAAAAAGATGAATTTTATTCC and GGATCC(BamHI)/AGCTTTAATAGTGTGCAGGGTCAGTCAG). PCR products were purified by PCR clean-up/Gel extraction kit (MACHEREY-NAGEL) and cloned into pBluescript SK (+). Plasmid DNA isolated from the transformed bacteria was sequenced.

### Antibodies

The primary and secondary antibodies used in this research were as follows: Anti-Rpsa (Santa Cruz, sc-376295), Anti-GAPDH (Proteintech, 60004-1-Ig), Anti-PEDF (Santa Cruz, sc-16956), Anti-His-Tag (Proteintech, 66005-1-Ig), Anti-βIII tubulin (Thermo Scientific Pierce, 2G10), Anti-DYKDDDDK epitope (FLAG) tag (Thermo Scientific Pierce, MA1-91878), Anti-HA (Thermo Scientific Pierce, 26183), Anti-E-Cadherin (Cell Signaling Technology, #3195), HRP-conjugated donkey-anti-mouse IgG (1:5000), HRP-conjugated donkey-anti-rabbit IgG (1:5000), HRP-conjugated donkey-anti-goat IgG (1:5000), Cy5-conjugated donkey-anti-goat IgG (1:200), TRITC-conjugated donkey-anti-mouse IgG (1:200).

### Histology and Immunohistochemistry

To analyze Rpsa expression, brains were dissected at E18.5 and fixed with paraformaldehyde/Phosphate-buffered saline overnight at 4°C. Fixed samples were cryo-protected by 25% sucrose/Phosphate-buffered saline for 48 hours at 4°C and then embedded with O.C.T. compound (Sakura). Cryo-sections (30 µm thickness) were cut by cryostat (Microm HM505 N) and air dried. Sections were rinsed three times in Tris-buffered saline and treated with 0.2% Triton X-100/Tris-buffered saline for 10 minutes at room temperature, followed by blocking for 30 minutes in 5% Bovine serum albumin/Phosphate-buffered saline supplemented with 0.25% Tween-20 to prevent nonspecific binding. Primary antibodies were diluted in blocking buffer, and sections were incubated in primary antibody overnight at 4°C. All secondary antibodies were diluted with blocking buffer and sections were incubated in secondary antibody for 30 minutes at room temperature. Sections were stained by 40,6-Diamidino-2-phenylindole, Dihydrochloride (DAPI, 600nM) and embedded with 90% glycerol made with Tris-buffered saline.

To analyze neuronal morphology, brains were dissected at postnatal day (P) 3 or P15 and fixed with paraformaldehyde/Phosphate-buffered saline overnight at 4°C. Fixed samples were processed as described above. Cryo-sections (60 µm thickness) were cut and stained by DAPI.

### *In Utero* Electroporation

*In utero* electroporation was performed as previously described (46, 53, 57). Briefly, pregnant dams were anesthetized with Avatin and the uterine horn was exposed. Plasmids (1 µg) were then injected into the lateral ventricle of E15.5 embryo brains. Electroporation at E15.5 will allow for transfection of cortical pyramidal neurons in layer 2/3 (26–28). Embryo heads were placed between electrodes with the positive anode angled toward the cortex and three electric pulses of 35 V were applied with 50 ms intervals by a CUY21 electroporator (Nepa GENE). Embryos were placed back into the uterus and allowed to develop uninterrupted until brain samples were collected at P3 or P15.

### Analysis of Neuronal Morphology

All images were obtained using a confocal microscope (Leica SP8) and the experimenter was blind to the phenotype at the time of imaging.

Apical Dendrite Orientation at P3 – The angle at which the apical dendrite extends with respect to the cortical plate at P3 was measured using the ImageJ software angle tool. A straight line perpendicular to the edge of the tissue was used as a reference point to start measuring the angle. Absolute values of the measured angles were used to calculate mean angles. Standard deviation projection images from z-projection photos produced from z-stack data were used for analysis.

Neuronal Morphology Analysis at P15 – ImageJ software measuring features were used to analyze neuronal morphology. Standard deviation projection images from z-projection photos produced from z-stack data were used for analysis. Sholl analysis was performed using the Sholl Analysis Plugin (Gosh Lab, UCSD) for ImageJ following the developer instructions.

Dendritic Spine Morphology Analysis at P15 – Spines were objectively characterized based on geometric characteristics. Spines longer than 1 µm were classified as filopodia, while spines shorter than 1 µm were classified as thin. Stubby and mushroom spines were classified based on morphological appearance. Spines with two heads were classified as branched. Photos of spines were taken at 63x with 5x zoom and standard deviation projection images were used for analysis.

### Subcellular Fractionation

The subcellular fractionation kit (NBP2-47659) from Novus Biologicals was used to fractionate COS-1 cells according to the protocol provided by the manufacturer.

### Pull-down Assay

Immunoprecipitation was performed as previously described (58). Briefly, transfected cells were lysed by NP-40 lysis buffer: 1M Tris pH 7.4, 5M NaCl, 0.5M EDTA, 20% NP-40, H_2_O. Supernatant was immunoprecipitated by anti-6XHis antibody conjugated Agarose beads (Santa Cruz). After being thoroughly washed by washing buffer, immunoprecipitated proteins were separated by SDS-PAGE and blotted with an anti-HA antibody.

### Cycloheximide Treatment

For the analysis of Rpsa turnover rate on the plasma membrane, cells were treated with cycloheximide (1 mg/ml) 48 hours after transfection. After 0, 3, 6, 12, 24, and 48 hours, sub-cellular fractionation was performed as described above to isolate the cytosolic and membrane fractions.

### Calcium Imaging of Live Brain Slices

Slice Preparation – Brain slices were prepared using a modified version of a previously described protocol (53). Briefly, male and female mice (approximately P30) were decapitated and brains were quickly removed and placed in ice cold high sucrose artificial cerebral spinal fluid (ACSF) solution for slicing. Brains were embedded in 4% low-melting agarose and coronal cortical slices were generated (300 µm) with a vibrating microtome (VTS1000 Leica Microsystems). Slices were next incubated for 60 minutes at 37°C in DMEM/F-12 imaging media without phenol red supplemented with 10% fetal bovine serum (FBS).

Imaging of Spontaneous Activity – The experimenter was blind to the phenotype at the time of the imaging. Slices were gently transferred into glass bottom 35mm dishes (MatTek) for imaging. A membrane was placed on top of the slices to reduce movement during imaging. Time-lapse live imaging was performed using an inverted fluorescent microscope (Ziess, Axio Observer Z1) with a 20x objective. Images were captured every 50 ms for 1 minute while the slices were maintained at 37°C with a stage top incubator (Zeiss).

Calcium Imaging Data Analysis – All cells included in the analysis were double-positive for GCamp6s and either pCAG-eCas9-tdTomato-U6-control-gRNA or pCAG-eCas9-tdTomato-U6-Rpsa-gRNA. All image analysis was performed using Zen 2 Pro analysis software (Ziess 2011). Circular regions of interest were placed on the cell soma. Baseline fluorescence (F_0_) was obtained by averaging the fluorescence intensity inside the region of interest immediately before beginning the time course imaging. Images were captured every 50ms for 1 minute. Fluorescence intensity for the time course was measured by averaging all pixels in the region of interest at each frame of the imaging (32). Percent change in fluorescence (ΔF/F_0_) was calculated as (Fmeasured- F_0_)/F_0_ for every frame of the time course.

### Electrophysiological Recordings

Slice Preparation – Mice (approximately P30) were decapitated and brains were quickly removed and placed in ice cold sucrose solution containing the following (in mM): 87 NaCl, 75 sucrose, 25 glucose, 25 NaHCO_3_, 1.25 NaH_2_PO_4_, 2.5 KCl, 0.25 CaCl_2_, and 3.5 MgSO_4_ for slicing. Coronal cortical slices were generated (300 µm) with a vibrating microtome (Leica Microsystems). Slices were next incubated for 30 minutes at 37°C in recording artificial cerebrospinal fluid (ACSF) containing the following (in mM): 111 NaCl, 3 KCl, 11 glucose, 25 NaHCO_3_, 1.3 MgSO_4_, 1.1 KH_2_PO_4_, and 2.5 CaCl_2_. Slices were then maintained at room-temperature in ACSF for at least 30 minutes before recording. Both slicing solutions and recording solutions were continuously aerated with 95%/5% CO_2_/O_2_.

Recordings – The experimenter was blind to the phenotype at the time of the recording. All recordings were performed at room temperature. Cortical neurons transfected with either the control CRISPR plasmid or the Rpsa CRISPR plasmid were visualized based on their fluorescence (tdTomato) with a 63x objective lens on a BX51WI scope (Olympus) using LED illumination (X-cite). Patch electrodes (Harvard Apparatus) were pulled to tip resistances of 5-8 M Ώ using a multi-stage puller (Sutter Instruments) and were filled with K-gluconate based intracellular solution, containing the following (in mM): 128 K-gluconate, 10 HEPES, 0.0001 CaCl_2_, 1 glucose, 4 NaCl, 5 ATP, and 0.3 GTP. Data was collected with a Multiclamp 700B amplifier (Molecular Devices) and Clampex software (pClamp9, Molecular Devices). Signals were digitized at 20 kHz and filtered at 4 kHz.

Resting membrane potential of the neuron was recorded in current-clamp mode immediately after breaking in. For passive and active membrane properties, current- and voltage-clamp protocols were performed as previously described(32). Briefly, input resistance was measured from a series of hyperpolarizing steps in voltage clamp using Clampfit. Time constant, tau, was calculated from the standard exponential fit of hyperpolarizing steps in current clamp. Capacitance was calculated from the input resistance and the tau measurements. Rheobase was defined as the lowest current step (in 10 pA increments) that evoked an action potential in the neuron.

### Statistical Analysis

Quantitative data were subjected to statistical analysis using SPSS (IBM Analytics), Prism (GraphPad Software) and MATLAB (MathWorks). The data were analyzed by two-tailed independent-samples t-tests, one-way ANOVAs or two-way ANOVA when appropriate. Values represented as mean ± S.E.M. Results were deemed statistically significant if the *p* value was <0.05. *, ** and *** indicate *p* <0.05, *p* <0.01 and *p* <0.001, respectively.

## Acknowledgements

We thank Dr. Bryan W. Luikart for his technical advice and support. We are also grateful to Drs. Peter Baas, Elias Spiliotis, Eric Olson, and Itzhak Fischer for their reading the manuscript and comments.

---

**Supplementary Figure 1. Validation of KD plasmids**

A) Western blots showing the efficacy of Rpsa KD, PEDF KD, and Itga6 KD done using 293T and COS-1 cells. B) Quantification of Western blots done using ImageJ shows Rpsa CRISPR efficiency is ~77%, PEDF shRNA efficiency is ~84%, and Itga6 shRNA efficiency is ~ 79%. Three biological/technical replicates were used (n = 3). Band intensity was quantified and normalized to GAPDH. C) Representative chromatograms for the target sequence of Rpsa CRISPR. D) Representative Rpsa gene alterations caused by Rpsa CRISPR. Numbers indicate amino acid position in Rpsa protein. Sequences in red font show the target of Rpsa gRNA and PAM sequence. E) Immunofluorescence staining of cortical primary neurons transfected with pCAG-eCas9-GFP-U6-Rpsa-gRNA plasmid upon plating and then fixed after 6 days in culture. Note that in the GFP-positive Rpsa KD neurons Rpsa staining is reduced compared to surrounding GFP-negative normal neurons. Scale bar = 25 µm. F) Quantification of Rpsa marker fluorescence intensity. Rpsa corrected total cell fluorescence (CTCF) is significantly decreased in neurons positive for Rpsa CRISPR (43,881.13 CTFC ± 5,156.70), as compared to untransfected Rpsa controls (85,504.51 CTFC ± 9,189.28), t(38) = 3.950, *p*<0.0005. G) Immunofluorescence staining of cortical primary neurons transfected with PEDF shRNA upon plating and then fixed after 6 days in culture. Note that in the GFP-positive PEDF KD neurons PEDF staining is reduced compared to surrounding GFP-negative normal neurons. Scale bar = 25 µm. H) Quantification of PEDF marker fluorescence intensity. PEDF CTCF is significantly decreased in neurons positive for PEDF shRNA (11,735.13 CTFC ± 2,019.42), as compared to untransfected PEDF controls (25,563.82 CTFC ± 6,629.78), t(33) = 2.142, *p*=0.040. I) Immunofluorescence staining of cortical primary neurons transfected with Itga6 shRNA upon plating and then fixed after 6 days in culture. Note that in the GFP-positive Itga6 KD neurons Itga6 staining is reduced compared to surrounding GFP-negative normal neurons. Scale bar = 25 µm. J) Quantification of Itga6 marker fluorescence intensity. Itga6 CTCF is significantly decreased in neurons positive for Itga6 shRNA (3,947.66 CTFC ± 819.61), as compared to untransfected Itga6 controls (21,478.55 CTFC ± 6,285.73), t(10) = 2.766, *p*=0.020.

**Supplementary Figure 2. Dendrite length at P3 after Rpsa KD**

*In utero* electroporation was performed at E15.5. One-way ANOVA determined there was no statistically significant difference in dendrite length between Rpsa CRISPR (27.68 ± 1.40) and control CRISPR (27.53 ± 1.56) at P3 (F(3,96) = 0.503, *p* = 0.681). For all groups, n=25 cells from 3 mice. Data are represented as mean ± SEM.

**Supplementary Figure 3. Layering of neurons following Rpsa, PEDF, and Itga6 KD**

A) Representative photos of pyramidal neurons expressing GFP or Venus at P15 after IUE at E15.5, scale bars = 100 µm. Rpsa and PEDF deficient neurons show mild defects in layering, with a broader distribution of somas in the CP as compared to control. Neurons deficient in Itga6 show no phenotype. B) One-way ANOVA showed a significant increase in mean distance of the soma from the edge of the tissue in Rpsa CRISPR (105.76 ± 3.99, *p*<0.0005) and PEDF shRNA groups (90.12 ± 4.66, *p*<0.0005) as compared to control CRISPR (41.34 ± 2.10) and scramble shRNA (43.26 ± 3.44), respectively (F(3,76) = 99.475, *p*<0.0005). For all groups, n=20 cells from 3 mice. Data are represented as mean ± SEM. C) One-way ANOVA showed a significant increase in mean width of soma distribution (distance between the cell that migrated the farthest and shortest distances) in Rpsa CRISPR (107.85 ± 7.68, n=11, *p*<0.0005) and PEDF shRNA groups (105.94 ± 5.75, n=10, *p*<0.0005) as compared to control CRISPR (45.02 ± 2.58, n=19) and scramble shRNA (50.44 ± 4.17, n=20), respectively (F(4,75) = 48.267, *p*<0.0005). For all groups, cells were analyzed from 3 mice. Data are represented as mean ± SEM.

**Supplementary Figure 4. Intrinsic properties of neurons in *ex vivo* brain slice preparation**

Control CRISPR and Rpsa CRISPR neurons display similar intrinsic membrane properties. A) Examples of a control CRISPR and Rpsa CRISPR neuron responding to a series of hyperpolarizing and depolarizing current steps from a holding potential of −65 mV. B) Quantification demonstrated that capacitance was significantly larger in the Rpsa CRISPR group. No other significant differences in membrane properties were found between the control CRISPR (n= 9 cells from 3 mice) and the KD group (n=7 cells from 2 mice).

**Supplementary Figure 5. Distribution of PEDF and Itga6 in the developing cortex**

Immunofluorescence staining of wild-type brain slices at E18.5 showing the spatial distribution of PEDF (A) and Itga6 (B). Scale bars = 50 µm (A) and 100 µm (B).

