## Supplemental Figures for "PEDF-Rpsa-Itga6 signaling regulates cortical neuronal morphogenesis"

**Supplemental Figure 1.** **Blazejewski et al.**

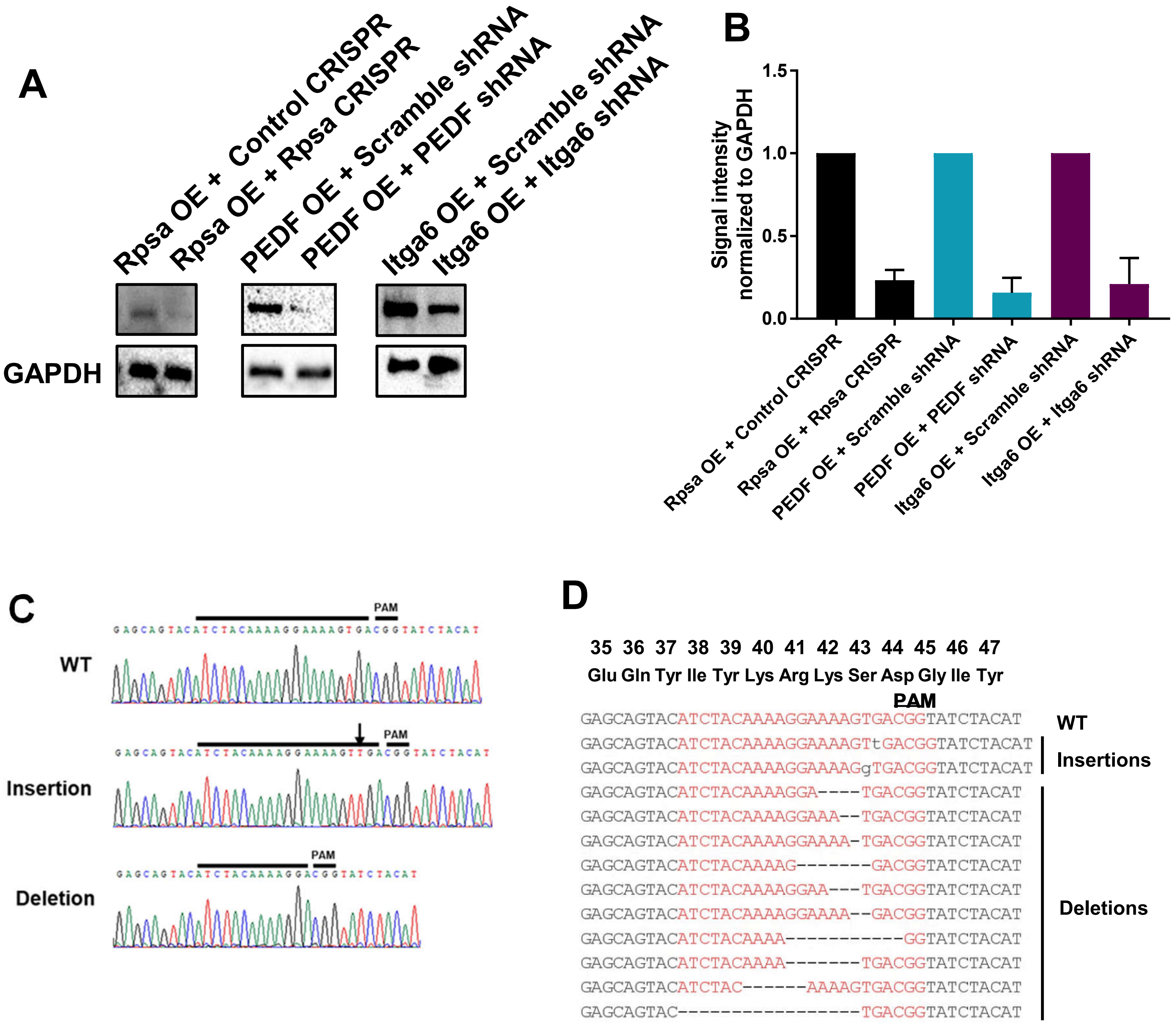

Supplemental Figure 1. continued

Blazejewski et al.

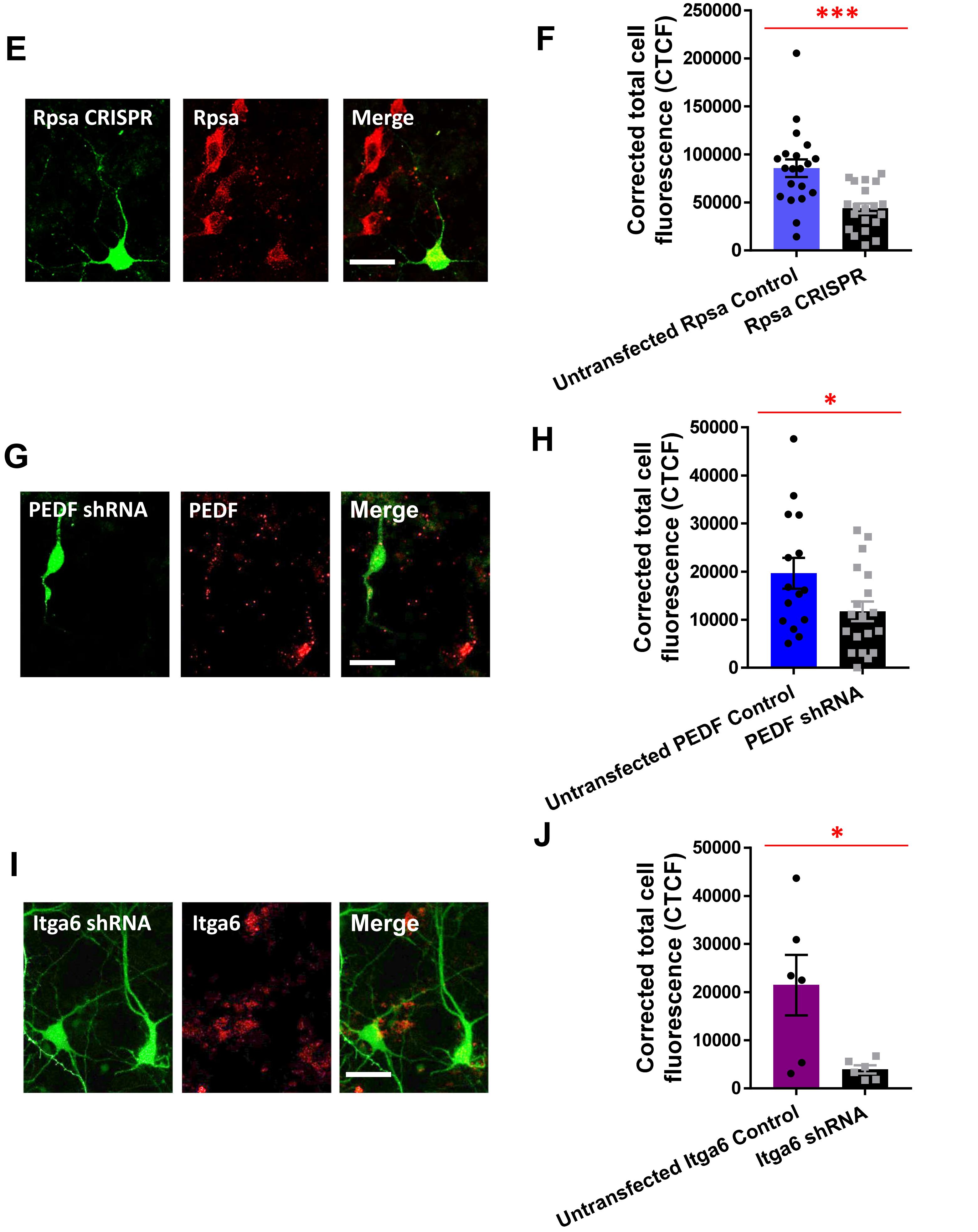

Supplemental Figure 2. Blazejewski et al.

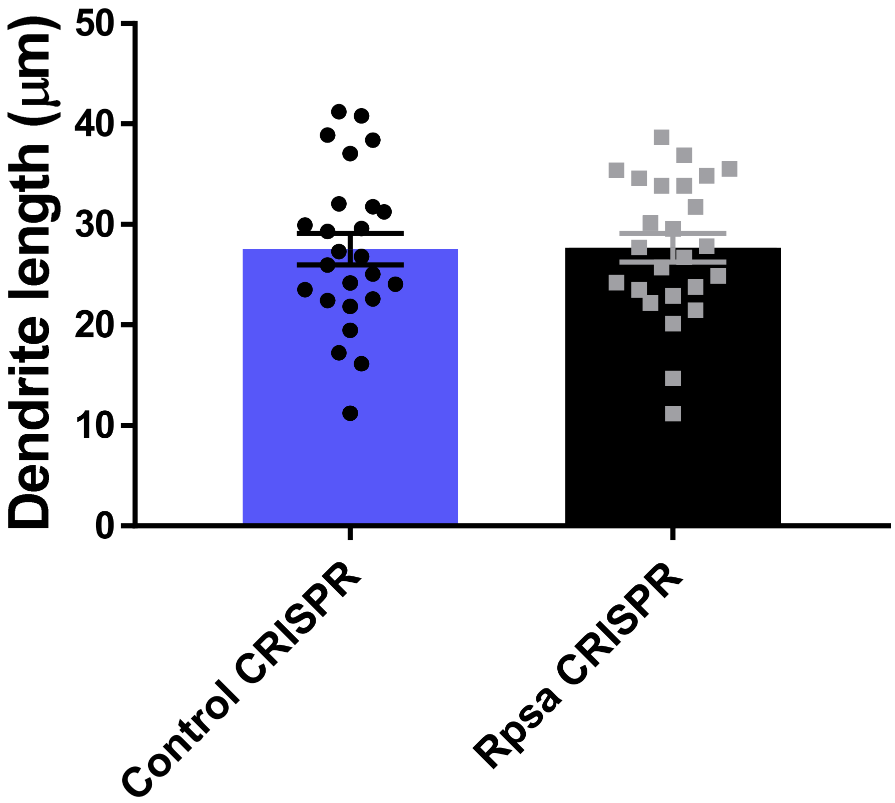

**Supplemental Figure 3.** **Blazejewski et al.**

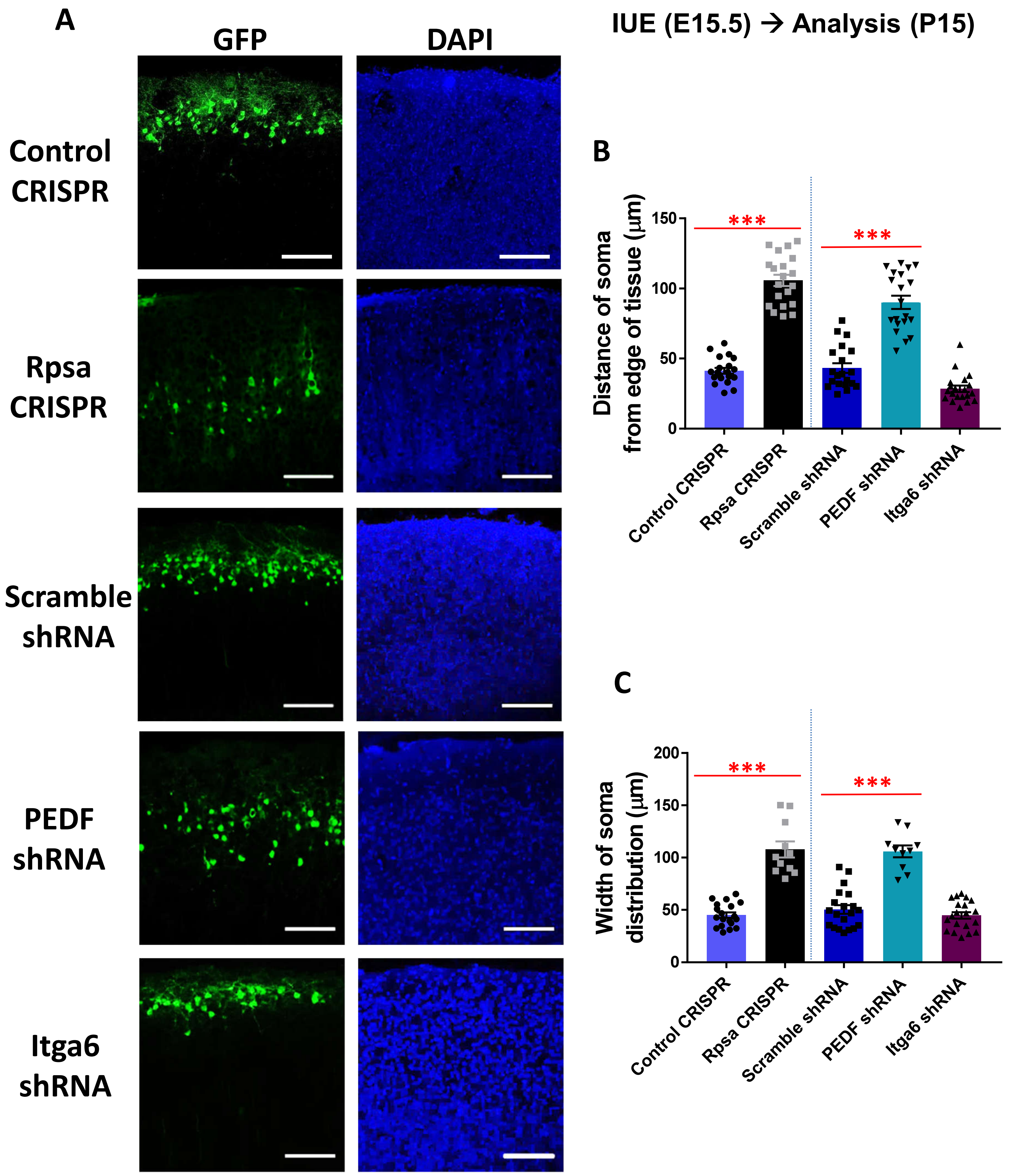

**Supplemental Figure 4.** **Blazejewski et al.**

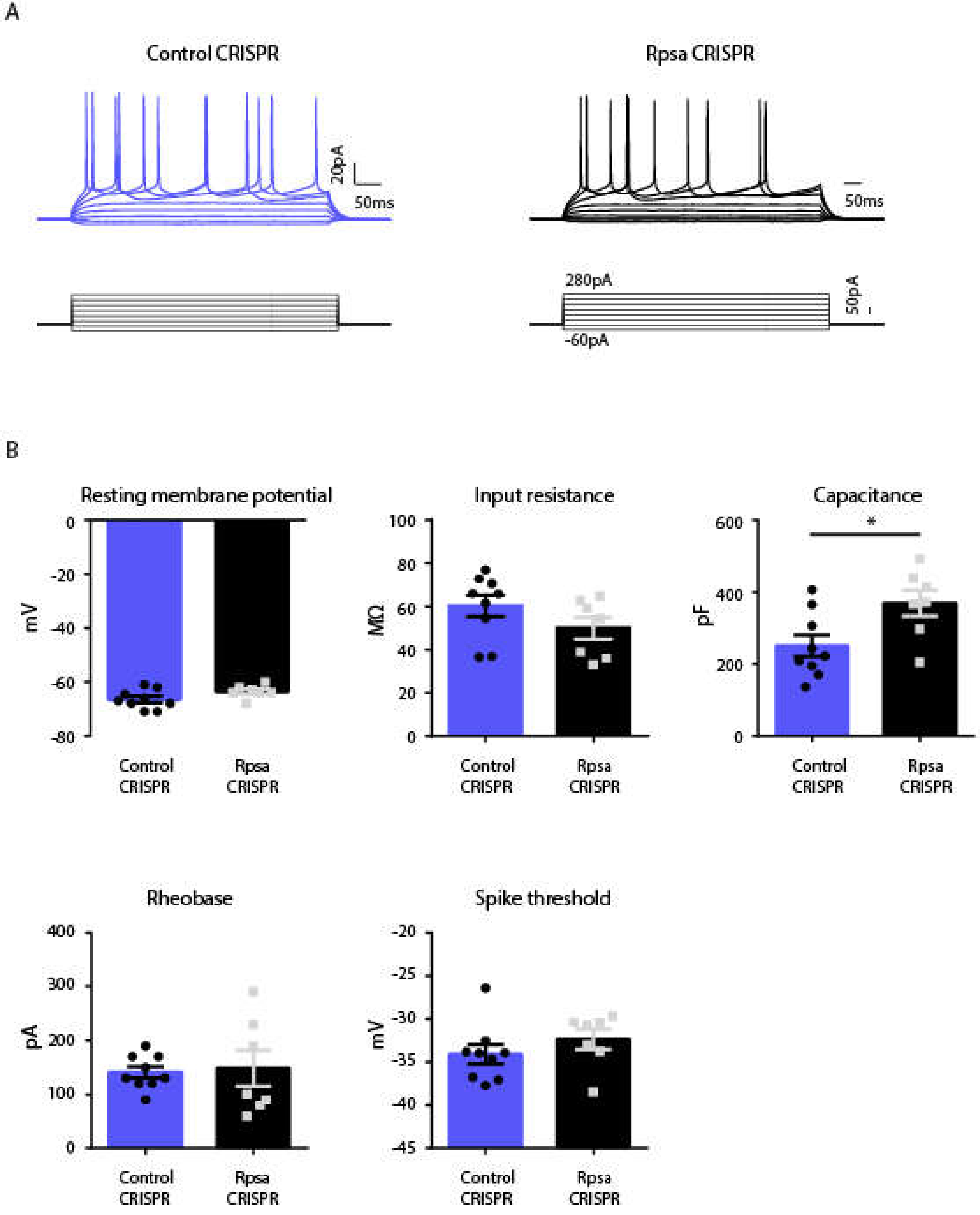

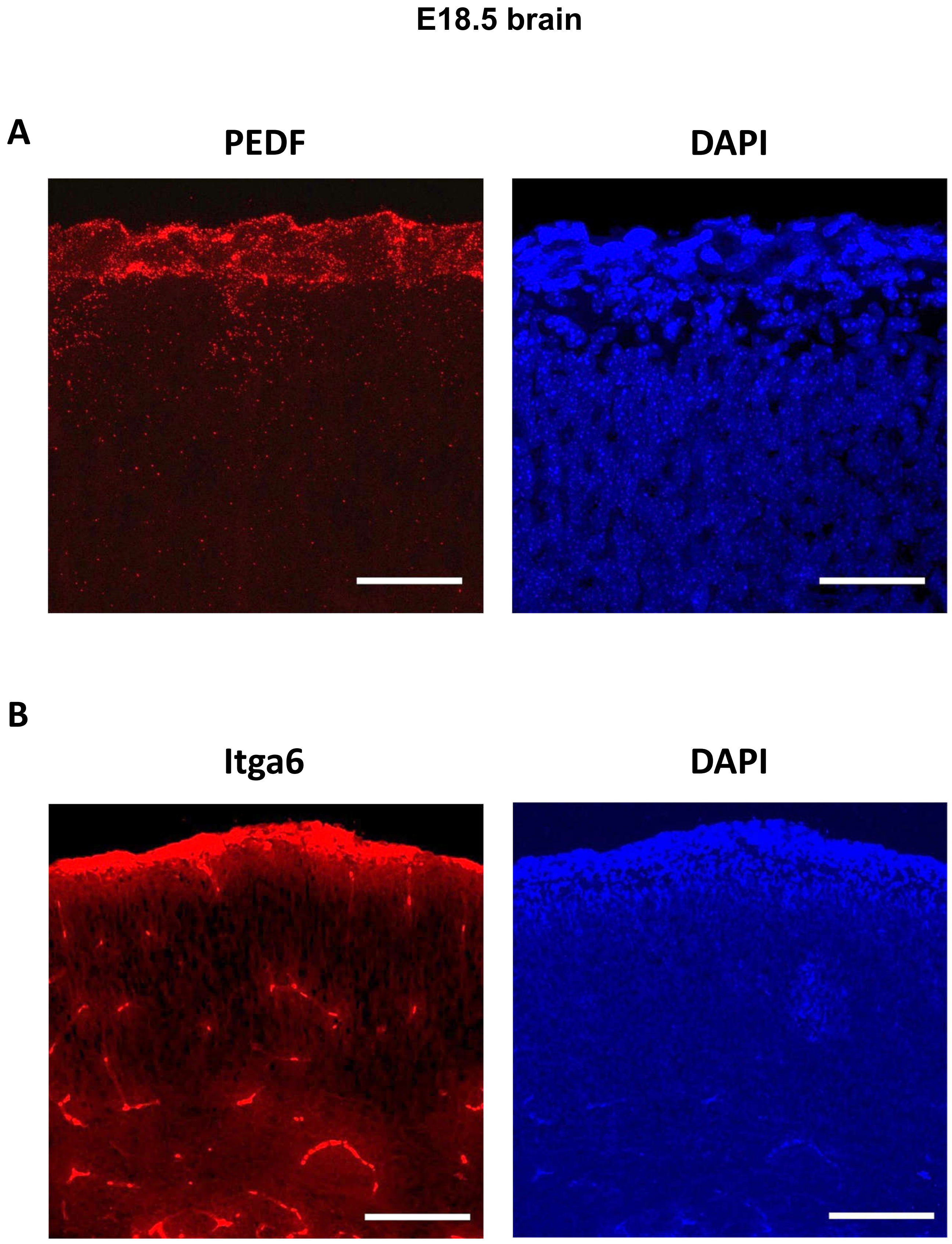
E18.5 brain
A
PEDF
DAPI
B
Itga6
DAPI
